# Noncoding regions underpin avian bill shape diversification at macroevolutionary scales

**DOI:** 10.1101/844951

**Authors:** Leeban Yusuf, Matthew C. Heatley, Joseph P.G. Palmer, Henry J. Barton, Christopher R. Cooney, Toni I. Gossmann

**Author notes:** **Correspondance:** (CRC) and (TIG).

## Abstract

Recent progress has been made in identifying genomic regions implicated in trait evolution on a microevolutionary scale in many species, but whether these are relevant over macroevolutionary time remains unclear. Here, we directly address this fundamental question using bird beak shape, a key evolutionary innovation linked to patterns of resource use, divergence and speciation, as a model trait. We integrate class-wide geometric-morphometric analyses with evolutionary sequence analyses of 10,322 protein coding genes as well as 229,001 genomic regions spanning 72 species. We identify 1,434 protein coding genes and 39,806 noncoding regions for which molecular rates were significantly related to rates of bill shape evolution. We show that homologs of the identified protein coding genes as well as genes in close proximity to the identified noncoding regions are involved in craniofacial embryo development in mammals. They are associated with embryonic stem cells pathways, including BMP and Wnt signalling, both of which have repeatedly been implicated in the morphological development of avian beaks. This suggests that identifying genotype-phenotype association on a genome wide scale over macroevolutionary time is feasible. While the coding and noncoding gene sets are associated with similar pathways, the actual genes are highly distinct, with significantly reduced overlap between them and bill-related phenotype associations specific to noncoding loci. Evidence for signatures of recent diversifying selection on our identified noncoding loci in Darwin finch populations further suggests that regulatory rather than coding changes are major drivers of morphological diversification over macroevolutionary times.

## Introduction

Disentangling the interplay between macroevolutionary trends and microevolutionary processes is fundamental to understand patterns of diversification over time. Key innovations, defined as traits that allow species to interact with environments in novel ways (Stroud and Losos 2016), are thought to play an important role determining macroevolutionary patterns of diversification, by allowing lineages to access and exploit new, previously inaccessible resources (Hunter 1998). In birds, evolutionary transitions in life-history traits and the emergence of *de-novo* innovations occurred rapidly alongside species and niche diversification (Balanoff et al. 2013; Xu et al. 2014). Understanding whether convergent molecular mechanisms underlie independent trait evolution in different organisms is a key question in biology (Manceau et al. 2010; Rosenblum et al. 2014; Lamichhaney et al. 2019). A variety of approaches to link molecular and phenotypic changes have been developed (O’Connor and Mundy 2009, 2013; Mayrose and Otto 2011; Levy Karin et al. 2017; Sharma et al. 2018; Hu et al. 2019) but these are generally restricted to relatively simple discretized phenotypic information (Prudent et al. 2016) and may not be easily applicable to more complex phenotypes on a genome-wide scale (Lartillot 2013).

A pertinent example of an important innovation is the evolution of the beak in modern birds. The avian bill is closely associated with species’ dietary and foraging niches and changes in beak shape are implicated in driving population divergence and speciation (Grant and Grant 1996; Bhullar et al. 2016). However, despite considerable effort, the genetic and developmental underpinnings of avian beak shape is still poorly understood, particularly at macroevolutionary scales. In the wake of the Cretaceous-Paleogene (K-Pg) mass extinction event, beak shape has been hypothesized to have evolved through a series of ontogenic stages (Bhullar et al. 2012, 2015), though the exact mechanism is yet to be established. Beak shape is comprised of separate morphological and developmental parameters, each of which is likely to be regulated by independent sets of transcriptional factors (Bhullar et al. 2015; Mallarino et al. 2011). Understanding how each of these morphological parameters evolved, how they are modulated, and how changes in such factors affect patterns of beak shape disparity across modern birds represents a significant unresolved challenge.

Several candidate genes linked to bird beak shape have been identified within populations or between recently diverged species. Among the earliest studies to identify genes implicated in beak shape evolution are comparative transcriptomic analyses in Darwin’s finches (Abzhanov et al. 2004, 2006) that found *BMP4*, a gene involved in the regulation of beak depth and width, and *CAM* (calmodulin), a gene putatively involved in beak length. Both genes were later identified to be partially-implicated in beak shape development (Mallarino et al. 2011). In addition, *ALX1*, a transcription factor involved in craniofacial development, and *HMGA2*, a gene associated with increases in beak size, were also identified in Darwin’s finches (Lamichhaney et al. 2015, 2018). In European populations of great tits, a collagen gene, *COL4A5*, putatively linked to beak length variation, was found to be under selection (Bosse et al. 2017a). Collectively, these findings illustrate (I) a complex genetic architecture for beak shape, (II) that genes implicated in beak shape may evolve under strong, detectable selective pressures in populations, and (III) that such genes are likely to be different across different avian taxonomic groups.

However, despite these clear predictions, no previous attempts have been made to identify genes that repeatedly play a role in beak shape evolution over broad evolutionary timescales. While previous studies have explored the genetic basis of other key avian traits (e.g. song, flight), such studies are typically targeted towards candidate genes, or incorporated clade-specific features (Whitney et al. 2014; Wirthlin et al. 2014; Machado et al. 2016; Sackton et al. 2019). Thus, relatively little is currently known about the genetics underpinning the macroevolution of beak shape. The current lack of insight connecting species or clade specific candidate genes to large scale evolutionary time may be explained by two main arguments. First, there is a growing consensus that large-effect genes (Fisher 1930) may not be as important for the evolution of complex traits as small-effect genes (Hill 2010; Rockman 2012; Boyle et al. 2017). This model of adaptation is well-supported by growing population genomic evidence, but does not explain candidate genes implicated in beak shape evolution with seemingly large-effects on beak morphology and speciation. Second, genes under strong long-term selective pressures may simply be difficult to detect due to confounding factors that obscure evolutionary signals. For example, selective pressures and demography vary over time, making the detection of clear signals of adaptive evolution and other evolutionary forces using sequence divergence approaches challenging. A third possibility is that the role of convergent evolution (Stern 2013; Manceau et al. 2010; Rosenblum et al. 2014; Lamichhaney et al. 2019) is limited if different genes are involved in morphological changes in different parts of the phylogeny.

Here, we utilize large-scale comparative genomic and phylogenetically reconstructed geometric-morphometric data to identify candidate loci that relate to macroevolutionary shifts in trait evolution. Specifically, we ask whether rates of bird beak shape evolution are explained by loci that experience long-term, repeated shifts in molecular rates across distantly-related avian taxa. To accomplish this, we designed an approach to detect loci persistently implicated in beak shape evolution across lineages by integrating morphological data into substitution rate models in a phylogenetic framework. We analysed protein coding genes as well as noncoding conserved regions from 72 bird species and combined them with morphological information from all major avian orders and families spanning >97% of avian genera (Cooney et al. 2017). Using this approach, we were able to link genetic and morphological diversification on a macroevolutionary scale.

## Results

Previous work has identified several genes and genomic regions that are under selection as likely species-specific drivers of bird beak shape evolution (Table S1). In order to identify genes that play a role in beak shape evolution beyond a lineage or species specific scale, we performed sequence divergence analyses on protein coding genes and avian-specific highly conserved elements, possibly regulatory, regions.

### Detecting protein coding genes repeatedly implicated in beak shape evolution

To test whether protein coding changes of the same protein are repeatedly implicated with beak shape morphological change across taxa, we designed an approach that incorporates estimates of morphological trait evolution into a branch model of sequence diversification. Specifically, we estimated sequence divergence using the ratio of non-synonymous substitutions to synonymous substitutions (*d*_N_/*d*_S_), which provides an indication of selection acting at the protein level. Our model assumes that the rate of molecular evolution (*d*_N_/*d*_S_) varies between predetermined types of branches, but not between sites in a protein, which is a reasonable restriction for computational reasons (Yang 1998; Yang and Nielsen 1998). We obtained estimates of rates of beak shape evolution based on geometric-morphometric data for all branches and grouped them into ranked bins according to their respective rates of beak shape evolution (Figure 1). If protein coding genes drive morphological change we hypothesised a positive correlation between ranked bins – where bins increased in rates of estimated phenotypic evolution incrementally – and estimates of *d*_N_/*d*_S_. For the 10,238 genes included in our analysis, we set up a branch model assuming different *d*_N_/*d*_S_ for each bin. Accompanying this, for each binned model, we estimated *d*_N_/*d*_S_ in a null model assuming no difference in *d*_N_/*d*_S_ between bins. A comparison between the binned model and the null model using a likelihood ratio test will reveal whether there is significant variation in the rate of protein change across our grouped branches.

**Figure 1:**
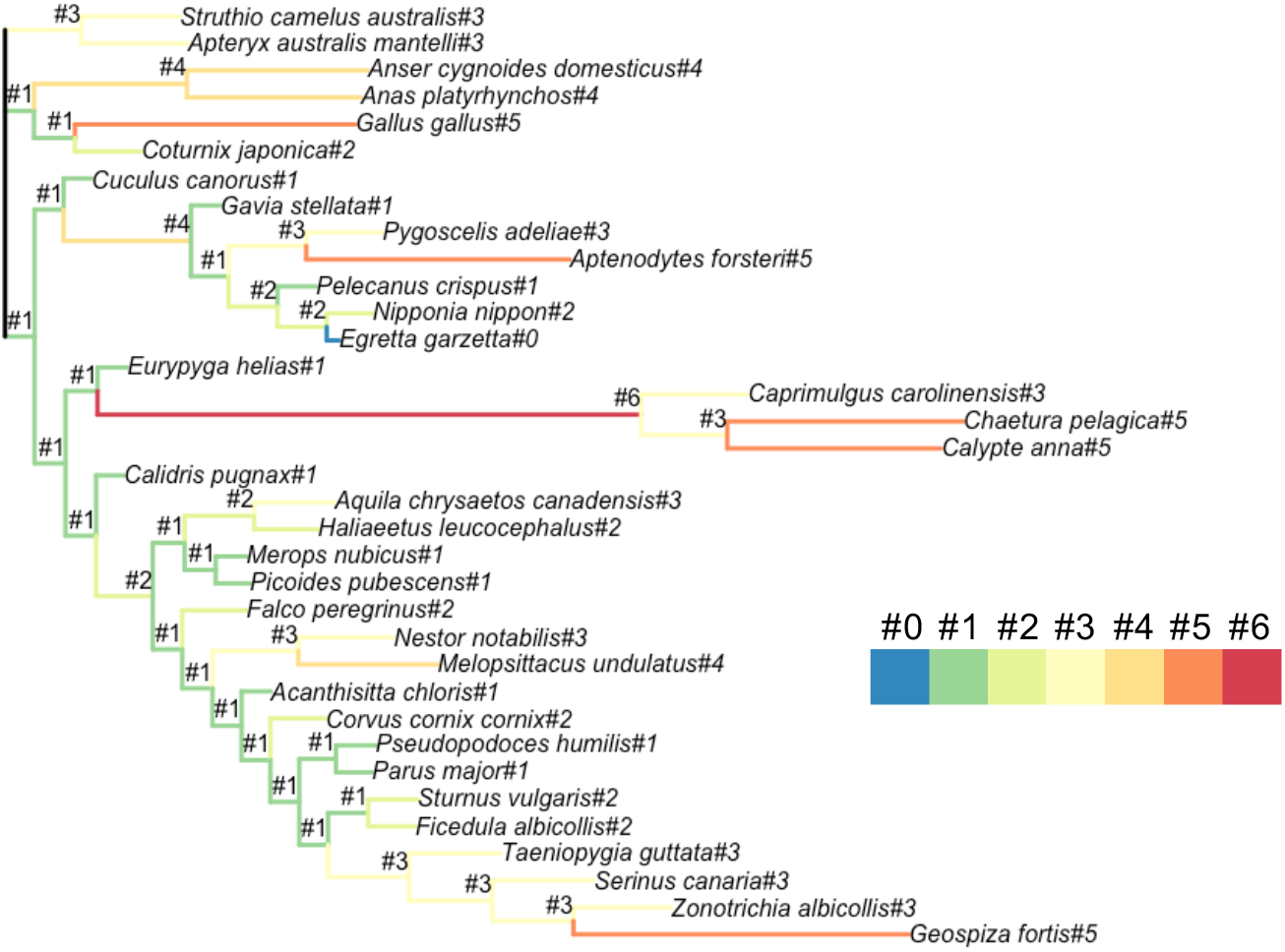
An example tree illustrating the grouping of branches according their beak shape morphological rates. The marked topology was then used as input for branch model in PAML (codeml for coding DNA and baseml for noncoding DNA). The maximum number of bins is eight for the coding gene set and 16 for the avian-specific highly conserved elements (ASHCE) set. Here, as an example, a binning with seven bins (#0 to #6) is shown.

We found that 1,434 (≈ 14%) genes had significant variation in their *d*_N_/*d*_S_ values across the grouped branches after correcting for multiple testing (e.g. significantly different likelihoods between the two models, FDR < 0.05). To determine putative functions of these genes, we performed phenotype ontology and pathway enrichment analyses using WebGestalt (Wang et al. 2017). Among the most enriched pathways are *Wnt Signalling pathway* and *ESC pluripotency pathways* (Figure 2A), both of which have been implicated in beak morphological development (Wu et al. 2004; Abzhanov et al. 2004; Merrill et al. 2008; Brugmann et al. 2010). Among the top phenotypic ontologies we find several ontology descriptions associated with skin as well as ectopic calcification and hydrocephalus (Table 1). We also used STRING (Szklarczyk et al. 2015), a comprehensive database combining different evidence channels for protein-protein interaction networks and functional enrichment analysis, to identify protein interaction partners of three proteins that have been previously identified as being associated with bird beak shape morphology independent of size effects (ALX1, BMP4 and CALM1, Table S1). ALX1, in contrast to BMP4 and CALM1, shows only two predicted interaction partners while the other two proteins are part of huge interaction networks (Figure S1). Altogether we identified 467 protein interaction partners across the three proteins and tested whether there is an enrichment of these in our dataset of 1,434 genes, which is indeed the case (Table 2, *χ*^2^ test, df=1, P=0.002).

**Figure 2:**
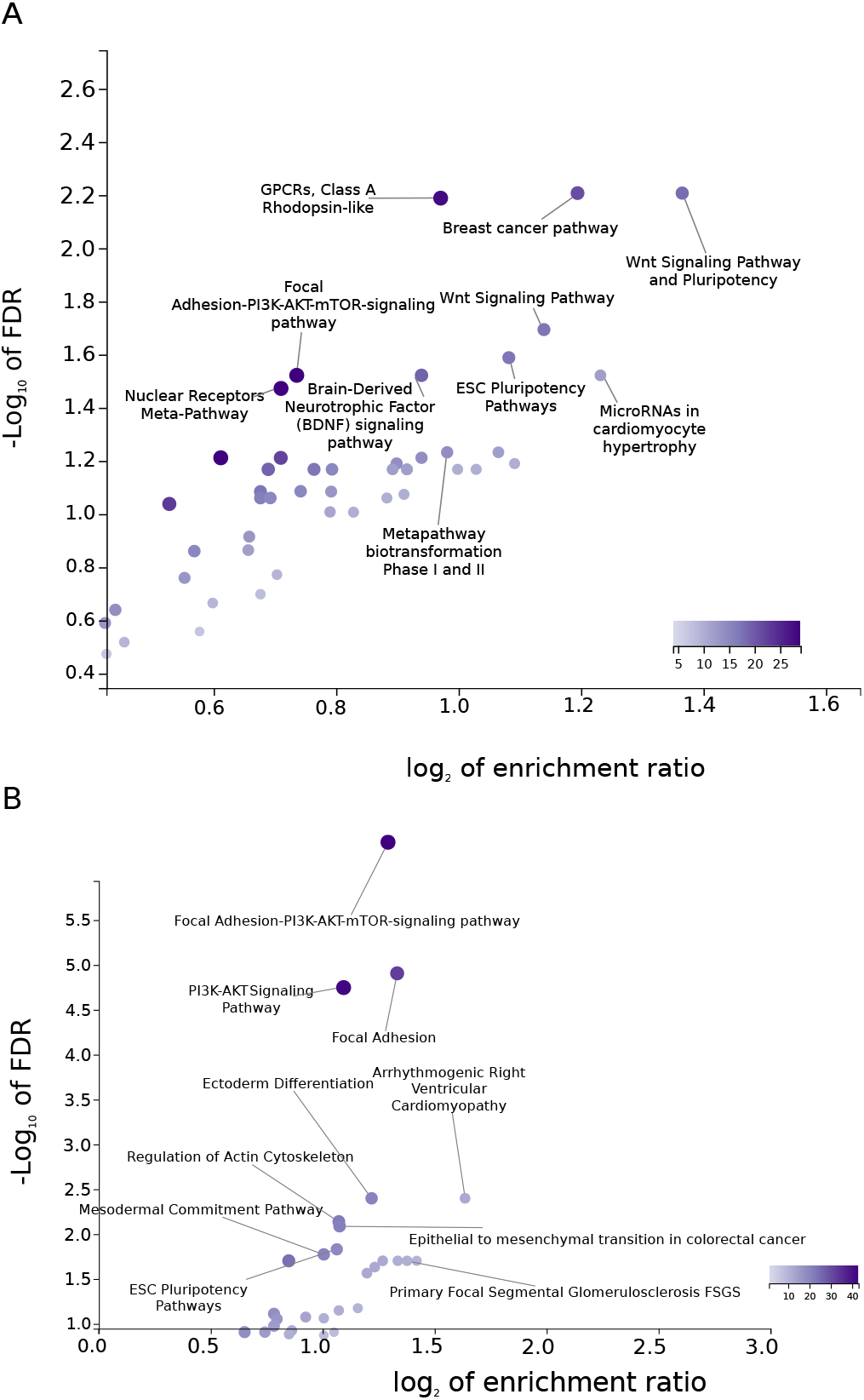
**Pathway enrichment analysis of (A) 1,434 protein coding genes and (B) 848 genes nearby avian conserved genomic regions** that show heterogeneity of substitution rates across branches that are grouped according to their beak shape morphological change rates. False discovery rate (FDR) and enrichment ratio stem from the pathway enrichment analysis in WebGestalt (Wang et al. 2017) using all analysed genes and human annotations, as these are the most comprehensive annotation databases to date. The color of the dots is denoted in the color scale and proportional to the category size, as definded by WebGestalt.

**Table 1:**
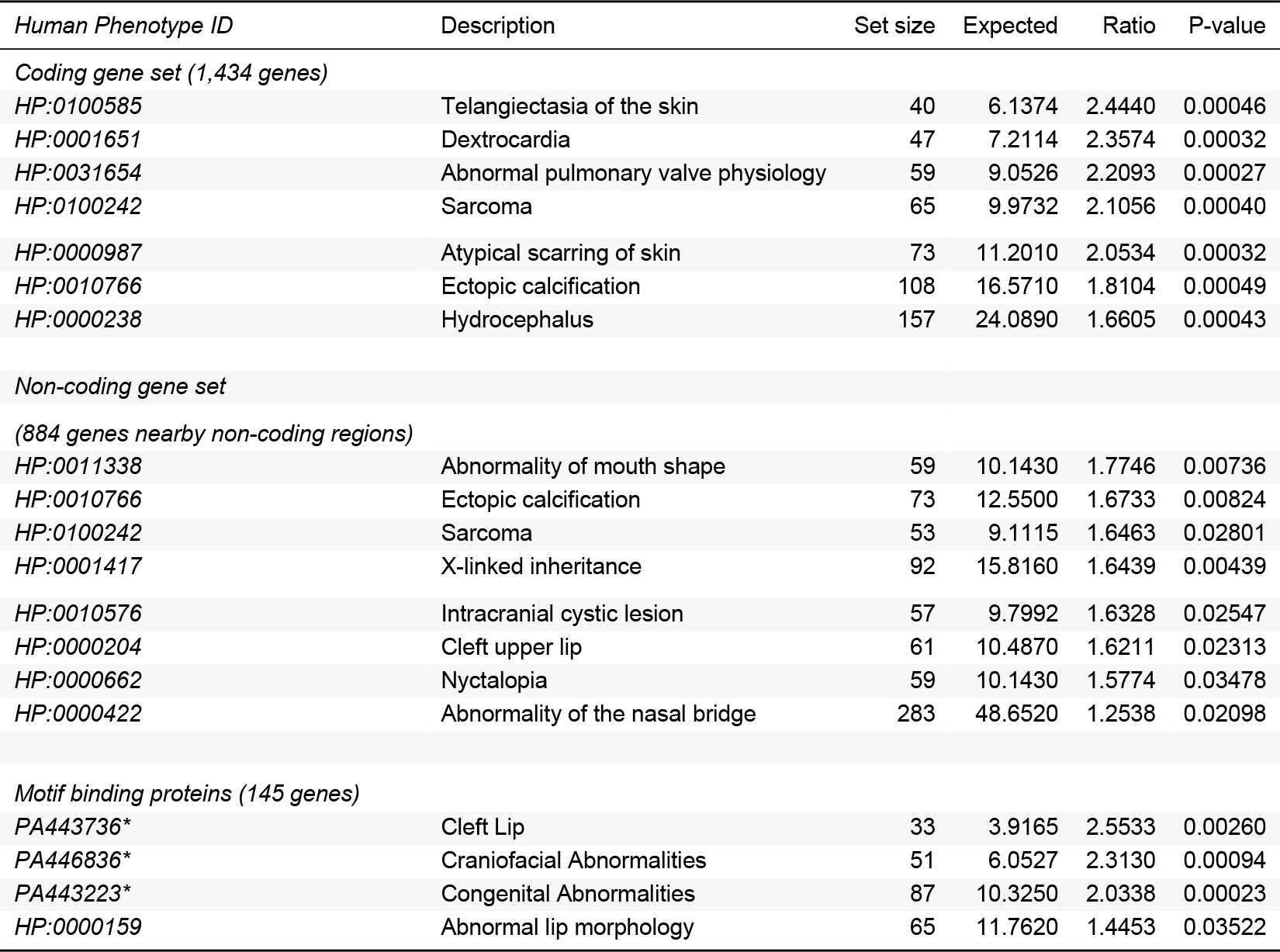
**Top phenotype ontology associations identified from the identified genomic loci**, coding genes, genes nearby noncoding regions and possible DNA binding proteins. * marked ontology terms are based on disease annotation database approach (GLAD4U).

| *Human Phenotype ID* | Description | Set size | Expected | Ratio | P-value |
| --- | --- | --- | --- | --- | --- |
| *Coding gene set (1,434 genes)* |  |  |  |  |  |
| *HP:0100585* | Telangiectasia of the skin | 40 | 6.1374 | 2.4440 | 0.00046 |
| *HP:0001651* | Dextrocardia | 47 | 7.2114 | 2.3574 | 0.00032 |
| *HP:0031654* | Abnormal pulmonary valve physiology | 59 | 9.0526 | 2.2093 | 0.00027 |
| *HP:0100242* | Sarcoma | 65 | 9.9732 | 2.1056 | 0.00040 |
| *HP:0000987* | Atypical scarring of skin | 73 | 11.2010 | 2.0534 | 0.00032 |
| *HP:0010766* | Ectopic calcification | 108 | 16.5710 | 1.8104 | 0.00049 |
| *HP:0000238* | Hydrocephalus | 157 | 24.0890 | 1.6605 | 0.00043 |
| *Non-coding gene set* |  |  |  |  |  |
| *(884 genes nearby non-coding regions)* |  |  |  |  |  |
| *HP:0011338* | Abnormality of mouth shape | 59 | 10.1430 | 1.7746 | 0.00736 |
| *HP:0010766* | Ectopic calcification | 73 | 12.5500 | 1.6733 | 0.00824 |
| *HP:0100242* | Sarcoma | 53 | 9.1115 | 1.6463 | 0.02801 |
| *HP:0001417* | X-linked inheritance | 92 | 15.8160 | 1.6439 | 0.00439 |
| *HP:0010576* | Intracranial cystic lesion | 57 | 9.7992 | 1.6328 | 0.02547 |
| *HP:0000204* | Cleft upper lip | 61 | 10.4870 | 1.6211 | 0.02313 |
| *HP:0000662* | Nyctalopia | 59 | 10.1430 | 1.5774 | 0.03478 |
| *HP:0000422* | Abnormality of the nasal bridge | 283 | 48.6520 | 1.2538 | 0.02098 |
| *Motif binding proteins (145 genes)* |  |  |  |  |  |
| *PA443736\** | Cleft Lip | 33 | 3.9165 | 2.5533 | 0.00260 |
| *PA446836\** | Craniofacial Abnormalities | 51 | 6.0527 | 2.3130 | 0.00094 |
| *PA443223\** | Congenital Abnormalities | 87 | 10.3250 | 2.0338 | 0.00023 |
| *HP:0000159* | Abnormal lip morphology | 65 | 11.7620 | 1.4453 | 0.03522 |

**Table 2:** Enrichment test for proteins identified as *d*_N_/*d*_S_ heterogeneous/homogeneous and interaction partners of BMP4/ALX1/CALM1. *d*_N_/*d*_S_ values were retrieved for groups of branches with similar beak shape morphological rates. The common interactome of BMP4/ALX1/CALM1 consists of 467 proteins, of which 256 were included in our gene analysis. Altogether, we identified 53 proteins of the BMP4/ALX1/CALM1 interactome that showed significant variation in their *d*_N_/*d*_S_ values (heterogeneous *d*_N_/*d*_S_).

| Protein category | $d_N/d_S$ | | Ratio |
| --- | --- | --- | --- |
| | heterogeneity | $d_N/d_S$ homogeneous | |
| BMP4/ALX1/CALM1<br>and interaction partners | 53 | 203 | 0.26 |
| Other proteins | 1381 | 8601 | 0.16 |
| P-value ($\chi^2$ 2 × 2 test,<br>df=1) | | | **0.002** |

We hypothesized a positive correlation between rates of molecular change and beak shape change, however after correcting for multiple testing no significant correlations were observed. This might be caused by limited power, e.g. due to short gene length or branch length, but generally suggests limited evidence for a simple relationship between the rate of molecular change in protein coding genes and morphological change.

### Detecting conserved noncoding regions implicated in beak shape evolution

To identify noncoding, possibly regulatory, regions that may be associated with beak shape morphological change over macroevolutionary time we analysed genomic regions based on avian conserved elements obtained from the chicken genome (Seki et al. 2017). Specifically, we obtained multiple sequence alignments of conserved regions from whole genome alignments comprising 72 bird genomes (Table S2) and grouped branches in up to 16 different categories using a *k*-means binning approach on branch specific morphological beak shape rate change (Cooney et al. 2017), a similar binning approach as for protein coding genes. Simulations show that 16 bins capture rate heterogeneity among branches very well at computationally feasible costs (Figure S2).

We successfully processed and analysed 229,001 conserved elements, of which 39,806 (≈ 17.4%) showed significant variation in their substitution rates after correcting for multiple testing (*χ*^2^-test, FDR<0.05). As we were interested to link potentially cis-regulatory elements to their target genes we restricted our analysis to conserved elements within or in close proximity to genes. Although the location of cis-regulatory elements is not fixed, they frequently occur in introns (Wittkopp and Kalay 2012) or close to the transcription start site, such as promoters and promoter-proximal elements (Butler and Kadonaga 2002; Andersson and Sandelin 2020). We extracted 884 genes that were overlapping or within 200bp distance (Piechota et al. 2010) of the identified regions in the chicken genome. To determine putative functions of these genes, we performed phenotype ontology and pathway enrichment analyses using WebGestalt (Wang et al. 2017). Among the most enriched pathways are *Ectoderm Differentiation, Mesodermal Commitment Pathway, Focal adhesion* and the *ESC Pluripotency pathways* (Figure 2B). The latter pathway set was also identified for the protein coding genes and represents an ensemble of pathways, including BMP and Wnt signalling, necessary for regulating pluripotency of embryonic stem cells (Okita and Yamanaka 2006). Phenotypic associations included “Abnormality of mouth shape and nasal bridge”, “cleft upper lip” and “nyctalopia” (Table 1).

We find that the 884 protein coding genes in cis to the identified genomic regions are overrepresented in a set of 511 genes involved in early craniofacial development in mice (Brunskill et al. 2014) (P=0.012, *χ*^2^-test, df=1, Table 3). To investigate whether there is any indicator for a biological meaningful relationship of the rate of molecular change and the rate of morphological change we focused on 2,644 out of 39,806 genomic regions (≈ 1.2% of all genomic regions) that individually showed a significant correlation (Kendall *τ*, P<0.05) between beak shape rates and substitution rate. We find that the over-representation for mice craniofacial genes is driven by a subset of 163 genes nearby the 2,644 regions (P=2.5×10^−5^, *χ*^2^-test), but not the 721 remaining genes (P=0.58, *χ*^2^-test, df=1). This suggests that the rate of molecular change in noncoding regions may be correlated to the rate of beak shape change (Table 3).

**Table 3:**
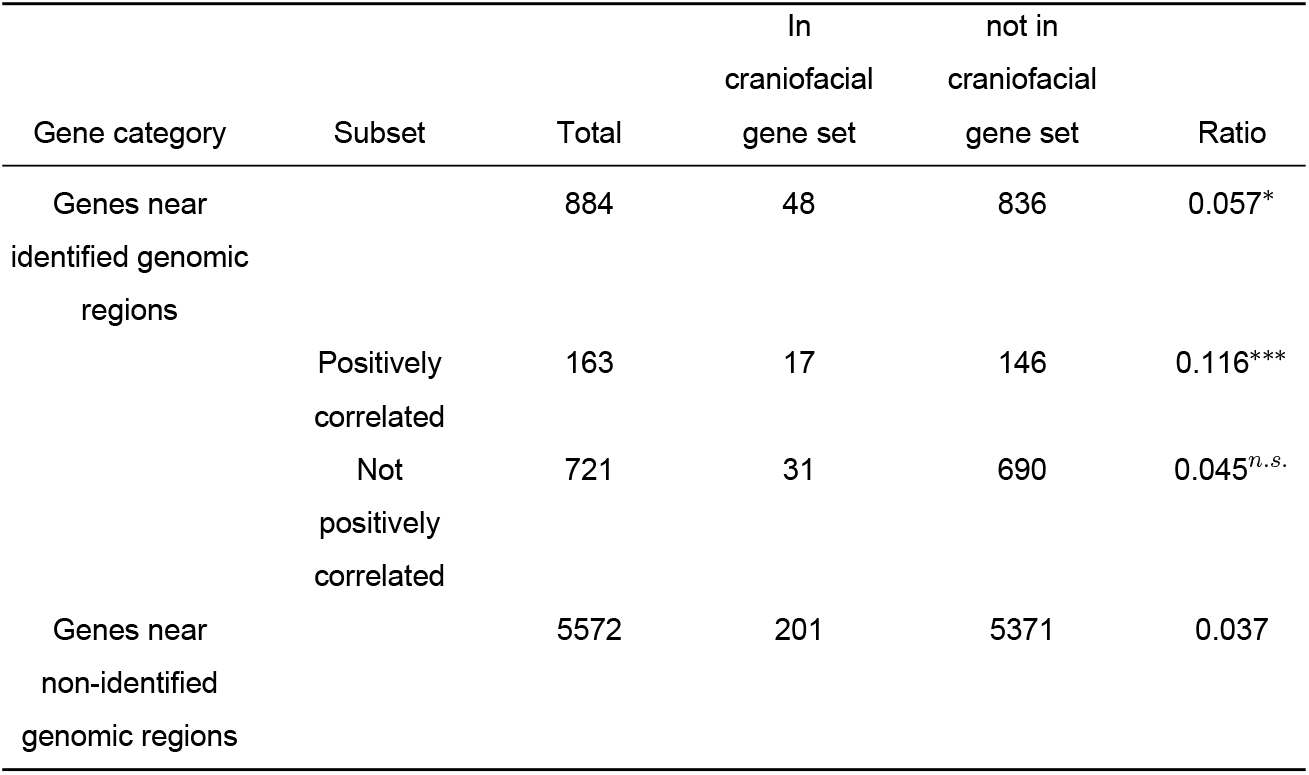
Enrichment tests for genes nearby genomic regions that show significant hetereogeneity in their substitution rates (heterogeneity was tested for grouped branches according to beak shape morphological change rates) versus a set of 511 known genes involved in craniofacial development in mice. Gene sets were further subset according to whether their was a significant correlation between morphological change of beak shape and substitution rates. P-values were obtained using a *χ*^2^ 2 × 2 test.

These previous analyses are most likely to identify the role of cis-regulatory elements because they focus on genes nearby to noncoding regions. Hence, we conducted a second strategy to gain further insights into the role of the identified noncoding regions as possible trans-acting elements. For this, we searched for short enriched motifs in the set of 39,806 genomic regions using DREME (Bailey 2011), and focused on the top 20 enriched motifs (Table S3). These motifs are potentially part of genomic regions that are targets of transcription factors.

To identify potential proteins binding to these motifs we used TOMTOM (Gupta et al. 2007) and obtained 145 potential annotated binding proteins, including GSC and SMAD proteins, both previously identified to be associated with beak shape morphological evolution (Parsons and Albertson 2009; Lamichhaney et al. 2015). To discern potential functions related to craniofacial features we conducted a phenotype enrichment analysis and identified “Abnormal lip morphology” as significant phenotype association and “lip and craniofacial abnormalities” as disease associated ontologies using a disease annotation database (Table 1).

### Genetic differentiation of the identified noncoding loci in Darwin’s finches

To test whether the identified noncoding loci may play a role in shaping beak morphology in a recent diversification we obtained polymorphism data from Darwin’s finch populations that either show a pointy or blunt beak phenotype (Lamichhaney et al. 2015). Using this dataset we find that our identified regions are characterised by patterns of linked selection that differ to a genomic control. Relative to genomic control regions, we find a stronger genetic differentiation between blunt and pointy phenotype populations (Figure 3A), as well as a higher overall genetic diversity at our identified loci (Figure 3B).

**Figure 3:**
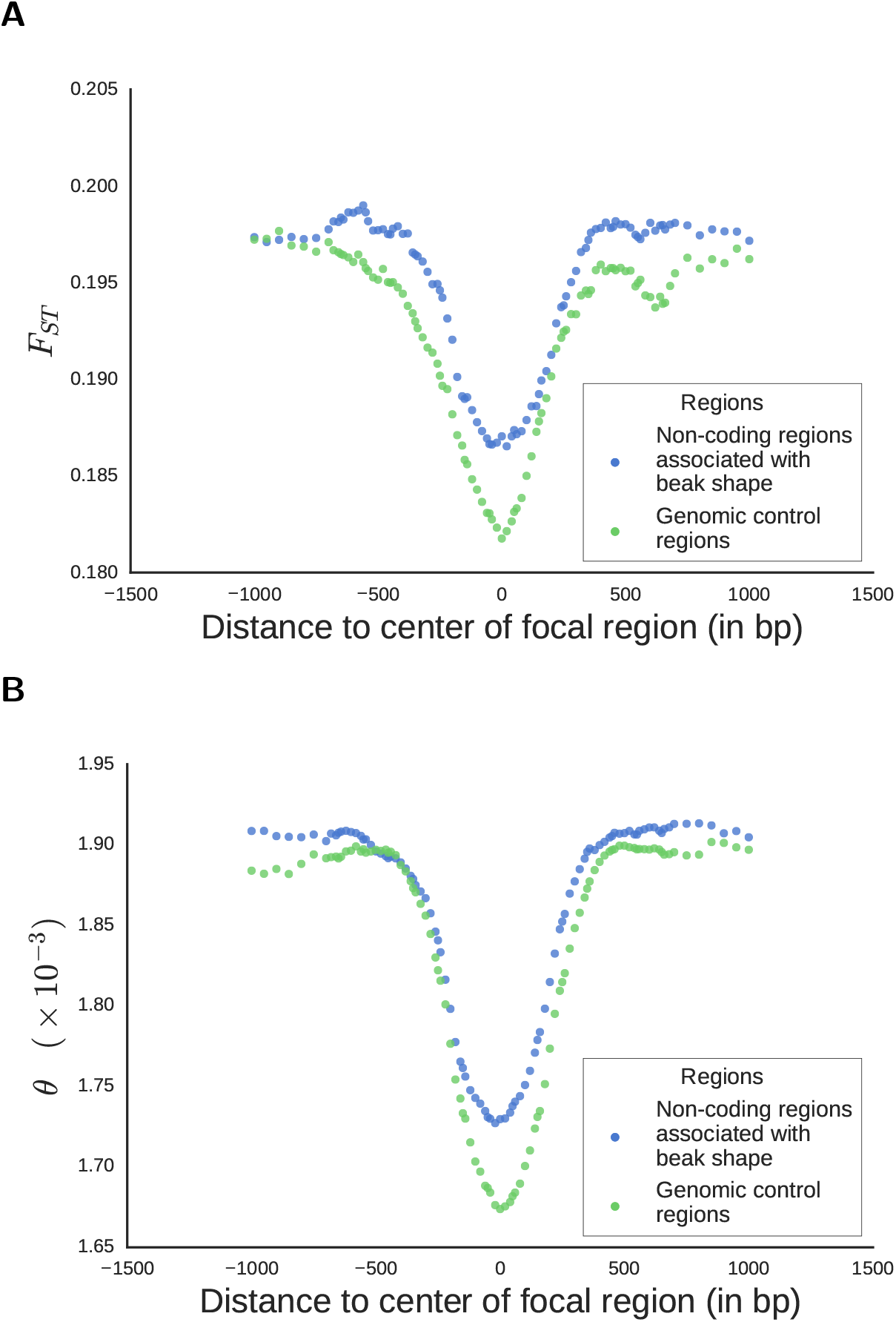
Identified noncoding genomic regions in a microevolutionary context in populations of Darwin’s finches. (A) Measures for genetic differentiation among populations, *F_ST_*, show contrasting genetic diversity in Darwin finch populations with blunt and pointy beaks, respectively. The identified loci associated with beak shape evolution over macroevolutionary time and nearby regions show a stronger differentiation relative to similar loci that are not asscoatiated with beak shape. (B) Total genetic diversity is higher for beak shape associated loci and nearby regions.

### Genes underlying evolutionary hotspots of beak shape divergence

Major evolutionary changes in beak shape may be concentrated within specific time periods and/or lineages (Cooney et al. 2017), and it is plausible that genes underlying these changes will show corresponding signatures of rapid evolution associated with such instances of ‘quantum evolution’ (Simpson 1944). We tested this prediction by identifying branches with the fastest-evolving rates of beak shape evolution according to trait evolution estimates derived from our morphological data. We selected three branches in our phylogeny with the most divergent beak shape evolution and refer to these branches as ‘hotspots’ (Figure S3). We conducted branch model tests (Yang et al. 1998; Yang 1998) for each of the three rapidly-evolving branches.

After accounting for multiple testing, we detected 36 genes with a signature of rapid evolution (*d*_N_/*d_S_* > 1, Figure 4). Although *d*_N_/*d*_S_>1 is indicative of rapid evolution, a formal significance test (versus a model with a fixed *d*_N_/*d*_S_=1 for the tested branch) suggests only for nine of our 36 identified genes a significant elevation of *d*_N_/*d*_S_ above one, indicative of positive selection (Figure 4). We identified BGLAP, a gene encoding osteocalcin, a highly-abundant, non-collagenous protein found in embryonic bone and involved in bone formation (Ducy et al. 1996; Raymond MH, Schutte BC, Torner JC, Burns TL 1999). Furthermore, we identified SOX5, a gene reported to have an assistive role in regulating embryonic cartilage formation (Lefebvre et al. 1998). In chickens, the expression of SOX5 and a duplication in the first intronic region of the gene is associated with the Pea-comb phenotype (Wright et al. 2009).

**Figure 4:**
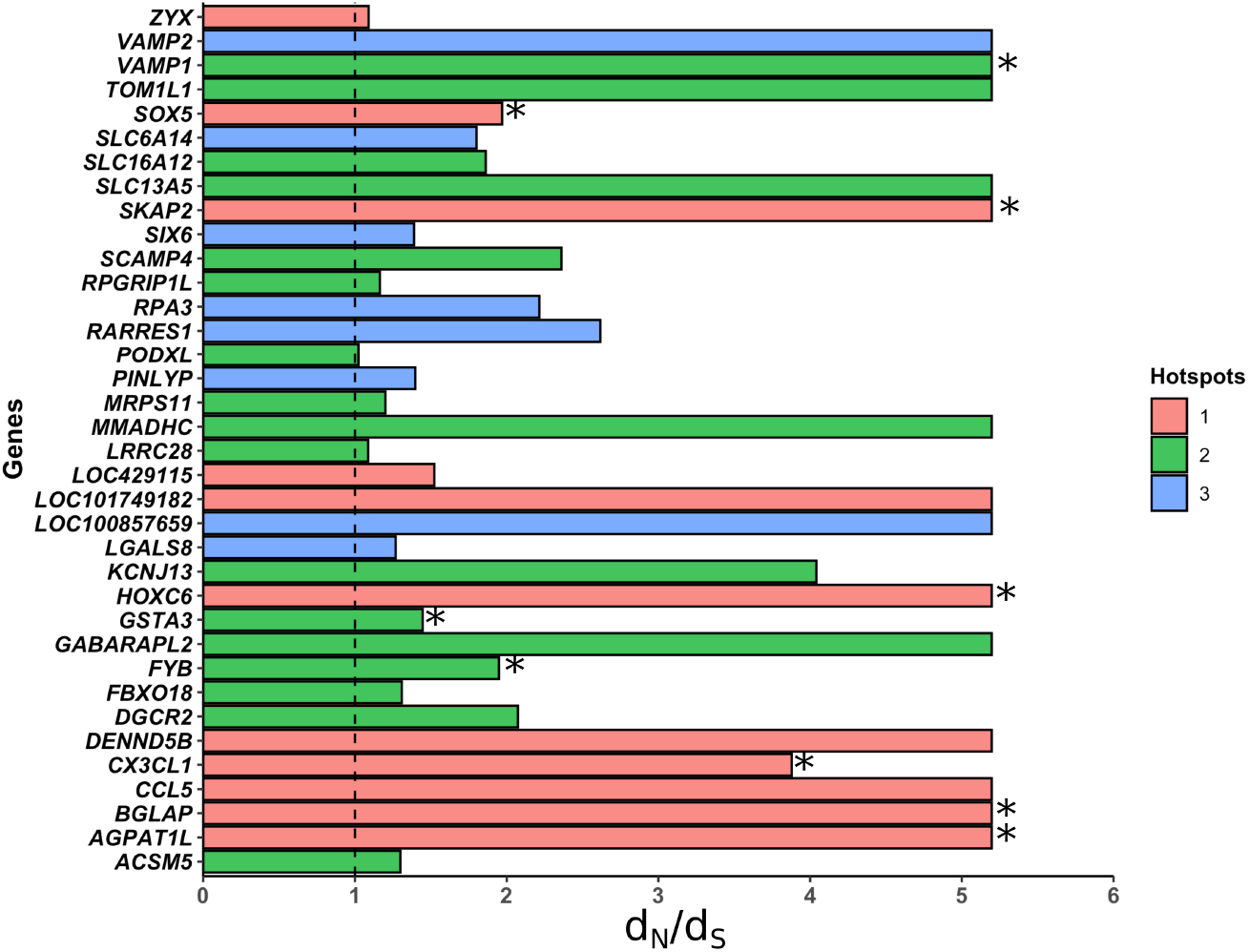
Elevated rates of protein evolution (*d*_N_/*d*_S_) associated with hotspots of beak shape morphological diversification. Shown are *d*_N_/*d*_S_ values for selected hotspot branches for 36 genes detected with *d*_N_/*d*_S_>1. Black dotted line formally indicates neutrality (*d*_N_/*d*_S_ = 1) and asteriks indicate genes for which the branch-specific estimate *d*_N_/*d*_S_ is significantly different from 1 (e.g. indicative of positive selection). Hotspots 1, 2 and 3 refer to the branches of the tree with the fastest, 2nd fastest and 3rd fastest rates of beak shape morphological change, respectively (Figure S3). For visualisation purposes are large *d*_N_/*d*_S_ values truncated at 5.2 (the estimate for SKAP2).

## Discussion

We developed a phylogenetic approach to identify genomic loci underlying the evolution of beak shape across macroevolutionary time and investigated genetic changes at coding and noncoding DNA across 72 bird species. Specifically, we asked whether loci that are repeatedly implicated in beak shape evolution across the bird phylogeny can be detected. By binning branches according to estimated rates of beak shape evolution, on the basis that phenotypic evolution is informative of genetic changes, we estimated rates of protein evolution across more than 10,000 genes, as well as rates of DNA substitutions for more than 200,000 avian-specific conserved regions, in a phylogenetic context.

### Protein coding genes associated with beak shape evolution across birds

For protein coding genes, we found significant variation in *d*_N_/*d*_S_ between binned branches in ≈ 14% of the genes tested, but we did not find a significant correlation between rates of phenotypic evolution and protein evolution for any gene. The binned model for coding DNA described in this study is not a formal test for positive selection, however a positive correlation between evolutionary rates and morphological genes could be indicative of repeated adaptive evolution of the same gene. Although we do not find evidence for this, some loci may have experienced shifts in *d*_N_/*d_S_* ratios repeatedly across distantly-related branches in association with beak shape morphological change. This relation may be explained by a number of different evolutionary forces, potentially acting independently or in tandem.

An association between *d*_N_/*d*_S_ and morphological changes may not be associated with adaptive events, but could also be explained by varying levels of genetic drift or purifying selection. For instance, relaxed purifying selection often occurs in response to environmental changes that weaken the effect of selection previously required to maintain a trait (Lahti et al. 2009). Furthermore, environments and therefore selective pressures are unlikely to remain stable over long evolutionary times. So far, only a few analyses have found experimental evidence of fluctuating selection acting on polymorphisms (Lynch 1987; O’Hara 2005). However, at a broader scale, models estimating the effects of fluctuating selection suggest a contribution to divergence similar to signatures of adaptive evolution (Huerta-Sanchez et al. 2008; Gossmann et al. 2014). Depending on the strength of fluctuating selection, or other types of varying selection intensities, it may account for the lack of strong, positive correlation coefficients reported in this study.

Although we might generally expect that morphological change in beak shape is positively correlated with *d*_N_/*d*_S_, an alternative scenario can explain a negative correlation. Adaptive mutations, because of their functional importance, are expected to experience strong purifying selection after their fixation (Kimura 1983). Functionally-important genes typically show signals of strong purifying selection (*d*_N_/*d*_S_ ≪ 1) and this is not conducive to a pattern of repeated increase in *d*_N_/*d*_S_ over distantly-related branches. Instead, a selective sweep would be followed by sustained reduction in *d*_N_/*d*_S_ through a prolonged period of intense purifying selection. We also did not identify genes with a significant negative correlations. We want to stress that further exploration of how adaptation occurs over macroevolutionary time, and the signals of selection left by ancient adaptive events is necessary to be able to fully elucidate our results. It may well be that our assumption of a positive correlation with beak shape rate does not hold because the role of convergent evolution is less pervasive, or that a rate analysis at coding sites does not have enough power as a measure of repeated directional positive selection.

### The effect of varying effective population size and life-history traits

Following the K-Pg extinction, modern birds experienced drastic reductions in body size, and with it, an increase in shorter generation time – this phenomenon is termed the Lilliput effect (Berv and Field 2018). Critically, reductions in body size and generation time have likely resulted in an increased *N_e_*, and alongside it, an increase in the efficacy of selection (Kimura 1983; Gossmann et al. 2010; Lanfear et al. 2014). This gradual decrease in body size and generation time, and with it, an increase in the neutral substitution rate, could account for an incremental decrease in *d*_N_/*d_S_* over time. So far, a number of studies have reported that a relationship exists between body mass and rates of molecular evolution in birds, with varying results (Weber et al. 2014; Nabholz et al. 2016; Botero-Castro et al. 2017; Figuet et al. 2017). In apparent contradiction with expectations of the neutral theory, several studies found that a decrease in body mass in birds did not result in decreased *d*_N_/*d*_S_ estimates (Lanfear et al. 2010; Nabholz et al. 2013; Weber et al. 2014; Bolívar et al. 2019). In fact, they found a weakly negative relationship between body mass and *d*_N_/*d*_S_, although similar studies report the opposite trend: *d*_N_/*d*_S_ in birds is positively correlated with body mass (Botero-Castro et al. 2017; Figuet et al. 2017). Indeed, mean and median correlation coefficient of *d*_N_/*d*_S_ with beak shape rate change is 0.047 and 0.048, respectively (significantly different from zero, P≪0.05, one sample *t*-test, n=1,434), for the 1,434 genes with significant heterogeneity in *d*_N_/*d*_S_, possibly suggesting a co-variation of beak shape change with other traits, such as body size.

Fluctuations in effective population size (*N_e_*), which are not taken into consideration by models of protein evolution, may affect interpretations of *d*_N_/*d*_S_. For example, fluctuations in *N_e_* may cause the fixation of neutral or slightly-deleterious mutations – in this case, this would mean the interpretation that *d*_N_/*d*_S_ > 1 is indicative of positive or diversifying selection may be erroneous. Strong shifts in *d*_N_/*d*_S_ may be driven by sudden changes in population size or genuine positive selection, and might obscure or oppose incremental increases in *d*_N_/*d*_S_ across bins (Bielawski et al. 2016; Jones et al. 2016). In our model, however, the effects of population size changes are partially negated by co-estimating parameters across branches. Unless specifically accounted for, substitution rate models do not consider the effect of non-equilibria processes that could affect *d*_N_/*d*_S_ estimates (Matsumoto et al. 2015). For example, GC-biased gene conversion - described as the preferential conversion of ‘A’ or ‘T’ alleles to ‘G’ or ‘C’ during recombination induced repair – has been shown to significantly affect estimates of substitution rates, in particular at synonymous sites in birds (Galtier et al. 2009; Weber et al. 2014; Boĺivar et al. 2016; Botero-Castro et al. 2017; Corcoran et al. 2017; Bolívar et al. 2019). However, while differences in the extent of GC biased gene conversion across genes are known, much less is known about its variation over time and incorporating such biases into large scale phylogenetic frameworks is far from trivial (Gossmann et al. 2018).

### Noncoding regions associated with beak shape morphology evolution across birds

Branch specific substitution rates of more than 39,000 avian-specific conserved regions are significantly associated with beak shape rates. We find more than 850 genes that are nearby these regions, possibly cis-regulatory factors, that show significant enrichment for craniofacial phenotypes in humans and mice. Unlike for protein coding regions we were unable to correct our substitution rate estimates for the effect of varying mutation rates (e.g. there is no counterpart for *d*_S_ as for coding regions). Due to special features of the avian karyotype, such as a stable recombinational and mutational landscape, it seems unlikely that variation in mutation rate can contribute to the patterns observed here. However, while inter-chromosomal re-arrangements are rare in birds, intra-chromosomal changes are more common and could lead to sudden changes in local mutation rates (Gossmann et al. 2018). Additionally, we restricted our analysis to noncoding regions that are specific to birds, or highly divergent relative to vertebrates (Seki et al. 2017). Whether anciently conserved elements, such as vertebrate specific regulatory regions (Lowe et al. 2015), may play a role in avian beak shape remains an open question. Equally, as the noncoding regions were identified based on the chicken genome, we lack those conserved regions that are absent from the chicken genome but present in other parts of the phylogeny.

More than 2,000 of the identified regions showed a significant correlation with binned rates of beak shape change and genes nearby these regions significantly overlap with genes involved in craniofacical development in mice (Table 3). The association of sequence divergence and trait divergence, along with a strong phenotypic enrichment, might suggest that the accumulation of neutral mutations at noncoding sites may play a pronounced role in bird beak diversification. This is because in our applied model we cannot distinguish between the action of selection and the accumulation of drift through fixations of new mutations (e.g. background variation in mutation rate). Hence, disentangling differences in the evolutionary pressures these regions experienced remains a major future challenge.

Some of the noncoding conserved genomic regions we identified may not be in physical proximity to a gene, e.g. many enhancers can be megabases away from the gene they regulate. Potentially, this could result from missing annotation for the *Gallus gallus* genome, or the fact that the genomic regions are trans-acting factors. Identifying the mechanisms underlying trans-acting factors is however very difficult to approach *in-vitro* as well as *in-silico*, and our approach to detect the role of trans-acting factors over macroevolutionary time is novel. We opted for an *in-silico* approach through motif enrichment and harvested vertebrate DNA binding protein databases to identify DNA binding proteins involved in beak diversification that go beyond a cis-acting role. We identified 145 possible DNA binding proteins using WebGestalt, including known transcription factors shown to be involved in beak development, that might be associated with beak shape diversification.

Although there is some overlap between the identified protein coding gene set and the noncoding gene set (Figure 5A), this is substantially less than expected by chance (P<0.005, *χ*^2^-test, df=1, protein coding genes versus genes near noncoding regions, Figure 5B). Indeed, based on the pathway and phenotype associations we note that the identified ontologies are different between the two datasets (Table 1). Although genes nearby noncoding regions are associated with facial and anatomical features, such as mouth shape, cleft and nasal abnormalities, the protein-coding phenotypes are mainly associated with dermal features. This suggests that the underlying evolutionary mechanisms of protein coding genes and noncoding, potentially regulatory, regions may be rather distinct in beak morphology evolution. However, as a common pattern, we identified that the ESC pathways are enriched in the coding and noncoding gene sets (Figure 5C). This further supports the notion that fundamental cellular pathways, such as BMP and Wnt signalling pathways, play a crucial role in the development of bird beaks and that this signal is detectable at a macroevolutionary scale.

**Figure 5:**
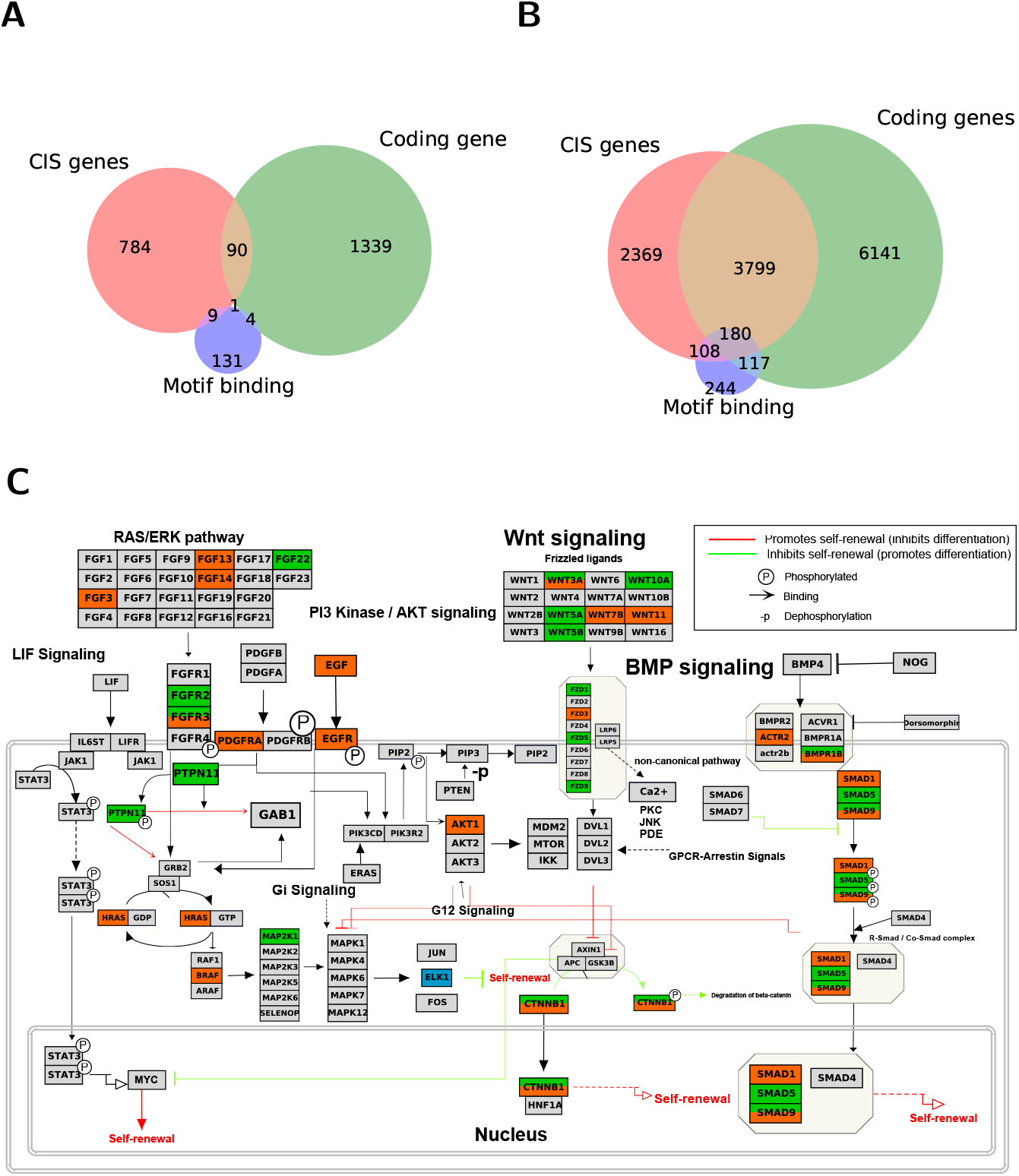
Comparison of beak shape associated gene sets derived from coding and noncoding genomic regions. (A) Overlap of the identified gene sets (B) Overlap of genes included in each dataset (background) (C) 32 Genes identified in our study occurring in the ESC pluripotent pathways, including BMP and Wnt signalling. Genes highlighted in green were detected in the protein coding analysis, while genes highlighted in orange were detected in the noncoding analysis. ELK1, labelled in blue, was detected as one of the transcription motif binding proteins. Coding genes denote all genes analysed for the protein coding gene analysis, CIS genes are genes in local proximity to analysed noncoding genomic regions, Motif binding are annotated proteins from the motif binding identification with TOMTOM (Gupta et al. 2007).

A pressing question remains as to whether these long-term associations are also reflected in selection at the micro-evolutionary level (Shultz and Sackton 2019). To test this we obtained data from Darwin’s finch populations that differ in their beak shape morphology (Lamichhaney et al. 2015). Our identified regions are characterised by pattern of linked selection that differ from genomic control regions with stronger genetic differentiation between blunt and pointy phenotype populations (Figure 3A), as well as a higher overall genetic diversity at our identified loci, suggestive of diversifying or partially relaxed purifying selection (Figure 3B). The signatures of selection are embedded in a genetic environment that shows local reduction of diversity due to strong purifying selection at these regions, typical of highly conserved regions. Sophisticated analyses of pinpointed genomic loci will be pivotal for future studies to disentangle the selective forces at these sites.

### Rapid genetic evolution in hotspots of beak shape evolution

In a second approach we focused on the findings of previous studies: beak shape changes are driven by different genes in specific branches. Applying this rationale, we identified rapidly-evolving lineages from comprehensive trait evolution analyses specifically focused on beak shape evolution (Cooney et al. 2017), and tested genes at these branches for accelerated rates of corresponding protein evolution. We identified 36 protein coding genes with branch specific signals of rapid evolution, with nine of them showing evidence for positive selection. These genes are putatively linked to branch and lineage-specific changes in beak morphology.

The most plausible candidates were detected in an internal branch leading to the evolution of the Strisores, a clade estimated to have diverged over 60 MYA, comprised of swifts, hummingbirds, nightjars and their allies (Hackett et al. 2008; Prum et al. 2015; Cooney et al. 2017). As well as distinctively short beaks evolving in swifts and nightjars, the divergence of hummingbirds is characterized by significant changes beak shape, body size and metabolism. This is supported by reported accelerated rates of evolutionary change in multiple cranial modules in Strisores (Felice and Goswami 2017). Together, these changes encapsulate adaptive shifts that have occurred in the Strisores clade.

An important candidate that may explain some of these changes is *BGLAP*, a gene encoding for osteocalcin, a ubiquitous protein found in bones and whose presence is critical for normal bone development (Ducy et al. 1996). Instead of direct involvement in bone production, osteocalcin regulates insulin expression and excretion, thereby regulating energy expenditure in muscle tissue, development of bone tissue and insulin sensitivity (Lee et al. 2007; Karsenty and Ferron 2012; Mera et al. 2016). Equally, as with *COL4A5*, a type IV collagen protein encoding gene, and *ALX1*, implicated in craniofacial development in Darwin’s finches, *BGLAP* may alternatively play a role in beak shape evolution (Lamichhaney et al. 2015; Bosse et al. 2017b).

Similarly, we identified *SOX5*, a gene previously associated with the evolution of craniofacial phenotypes in chickens, to be under putative positive selection in Strisores (Wright et al. 2009). Specifically, pea-comb development is associated with ectopic expression of *SOX5* caused by copy-number variation at the first intron of *SOX5* (Wright et al. 2009). This is independently corroborated by strong expression patterns of *SOX5* in the brain tissue – a possible proxy for craniofacial tissue, which is not included in expression profiles – of chickens (Merkin et al. 2012). Beyond the pea-comb phenotype, SOX5 is an essential transcription factor that acts to regulate chondrogenesis by enhancing a type-2 collage protein (COL2A1) and promotes the differentiation of chondrocytes. Critically, the expression pattern of *COL2A1* in the pre-nasal cartilage, an important morphological module of beaks and their shape(s), explains beak shape differences between medium and large ground finches during the 27th embryonic stage of development (Mallarino et al. 2011). Therefore, we suspect that *SOX5* may be important in explaining beak shape changes in swifts, hummingbirds and nightjars.

### Beak shape as a proxy for trait diversification

A key principle of adaptive radiation theory is that diversification of species is associated with ecological and morphological diversity (Schluter 2000). In birds, the evolution of morphological changes tends to coincide with speciation events, with some discontinuities, particularly early on in avian evolution (Foote 1997; Ricklefs 2004; Hughes et al. 2013; Mcentee et al. 2018). Here, we focus particularly on beak shape evolution because of its putative importance as a key ecomorphological trait and its link to speciation, demonstrated by long term trends and direct ecological evidence in Darwin’s finches (Cooney et al. 2017; Ricklefs 2004; Mcentee et al. 2018; Lamichhaney et al. 2018; Han et al. 2017; Grant and Grant 2009; Podos 2001; Huber and Podos 2006). However, evidently, many of the genes detected in this study are not associated with beak shape according to their putative functions. There are two explanations for this: First, some of the identified genes are pleiotropic in character and second, their functions are associated with traits that co-vary with beak shape evolution. We suspect that, alongside strong candidates for beak shape, we have detected genes implicated in a range of adaptive changes that have allowed species to diversify into different ecological niches. Here, estimates of beak shape evolution taken from Cooney et al. (2017) may have acted to identify branches with the fastest rate of phenotypic evolution rather than beak shape evolution specifically. This may be of particular relevance for the identification of genomic loci underlying beak shape diversification hotspots.

In summary, we were able to identify genomic loci associated with beak shape morphological evolution over macroevolutionary time by combining morphometric analyses with genomic data. For both, coding and noncoding regions, less than 20% of the tested loci show significant variation in their molecular rates, and most of the tested loci in this study are genetically very conserved on a macroevolutionary scale and hence cannot provide a genetic explanation for the observed phenotypic variation in beak shape across species. We show that homologs of identified protein coding genes as well as genes in close proximity to identified noncoding regions, are involved in craniofacial embryo development in mammals and pinpoint two associated pathways, BMP and Wnt signalling, illustrating that changes in coding as well as noncoding DNA facilitate phenotypic evolution of avian beak shape. The identified coding and noncoding loci are highly distinct, with significantly reduced overlap between them and fundamentally different phenotype associations. At present, the selective forces that contribute to patterns of genetic and morphological diversification remain difficult to pinpoint. However, as genomic and morphological data continue to accumulate, our framework offers a potentially powerful approach to further disentangling the interplay of selection and drift responsible for driving the diversification of complex phenotypic traits at macroevolutionary scales.

## Methods

### Multiple sequence alignments for protein coding genes

We used genomes of 57 bird species with high quality annotations from NCBI RefSeq (O’Leary et al. 2016) (Table S2). First, 12,013 orthologous protein coding genes were retrieved using RefSeq and HGNC gene identifiers, alongside reciprocal BLAST approaches based on three focal species, chicken, great tit and zebra finch - three of the best annotated high quality bird genomes available to date (Li et al. 2003; Östlund et al. 2009; Afanasyeva et al. 2018). We then performed a first set of alignment runs using PRANK (Löytynoja and Goldman 2008). To ensure the quality of these sequence alignments, we applied a customised pipeline including multiple alignment steps and quality filters. Details are described in the Supplemental Methods.

### Avian specific highly conserved regions (ASHCE) alignments derived from multispecies whole genome alignments

To estimate substitution rates for noncoding conserved elements across the bird phylogeny, we obtained whole genome information from NCBI for 72 bird species including the 57 bird species using in the coding DNA analysis. To generate a multispecies whole genome alignment we aligned the 72 avian reference genomes (as of 15/2/2017) against version 3 (galGal3) of the chicken (*G. gallus*) genome (version: galGal3 available from: ftp://ftp.ensembl.org/pub/release-54/fasta/gallus_gallus/dna/) using the MUL-TIZ package (Blanchette et al. 2004). Alignments were performed per chromosome following a pipeline published earlier (Corcoran et al. 2017). A list of query species, genome versions used and download locations can be found in Table S2. We used avian-specific highly conserved elements (ASHCE) from Seki et al. (2017). They used whole-genome alignments for 48 avian and 9 non-avian vertebrate species spanning reptile, mammal, amphibian and fish to obtain 265,984 ASHCEs. We were able to prepare 229,001 (86% of the total number of ASHCE) high-quality alignments as input for the analysis with baseml. For this target ASHCE regions were intersected with the whole genome alignments using BEDTools (v2.27.0) and FASTA files were created using customised scripts. Spurious and poorly aligned sequences were automatically removed using trimAl v1.4 (Capella-Gutierrez et al. 2009).

### Rates of morphological beak shape evolution

Information on beak shape evolution was extracted from a recent study (Cooney et al. 2017) that quantified patterns of beak shape evolution across 2,028 species (>97% extant avian genera) covering the entire breadth of the avian clade. Briefly, this study used geometric morphometric data based on 3-D scans of museum specimens and multivariate rate heterogeneous models of trait evolution (Venditti et al. 2011) to estimate rates of beak shape evolution for all major branches in the avian phylogeny.

Based on this information, we hypothesised that branches found to have experienced rapid beak shape evolution should also experience faster evolutionary change at the protein or genomic level. To test this, we split our evolutionary analyses into two, discrete approaches. First, for the detection of genes and genomic regions that have recurring effects on beak shape variation across multiple branches, we devised a binned approach. Second, for the detection of genes undergoing positive selection at branches that show rapid morphological change, we designed a hotspot approach.

### Binned branch approach for the detection of large-effect genes and regulatory regions

To detect genes that may be undergoing repeated periods of rapid, possibly adaptive, evolution across multiple lineages, we grouped branches in each alignment phylogeny according to their rates of morphological evolution using *k*-means binning (Lloyd 1982). Here, we opted for up to eight (coding) and 16 (ASCHE) bins, respectively, to enable robust statistical analysis but still reasonable computational time for the substitution rate analysis. To phylogenetically link the genetic data to the morphological data we relied on the Hackett et al. backbone (Hackett et al. 2008), hence we did not account for phylogenetic heterogeneity among genes and possible gene-tree species tree discordance. Branches were grouped incrementally based on rates of trait evolution using a *k*-means binning approach, with the first bin representing branches with the slowest rates of morphological evolution, and the last bin representing branches with the fastest rates of morphological evolution (Figure 1). We assumed that genes involved in beak shape evolution would experience evolutionary rate change at the protein level (*d*_N_/*d*_S_) proportional to their respective rate of morphological evolution. Theoretically, we hypothesize that genes important in beak shape evolution across many branches would show a strong positive correlation.

In our analysis, we tested this using a branch model which assumes different substitution rates (*d*_N_/*d*_S_) across different, pre-defined, branches in a phylogeny. Critically, the branch model may be useful in the detection of adaptive evolution occurring on particular branches (Yang et al. 1998; Yang 1998). Furthermore, we selected the branch model due to computational efficiency; the branch-site model and free-ratio model was deemed computationally intractable for a phylogeny of up to 57 species. Branches in each alignment’s phylogeny were marked according to their respective bins (typically ranging from 1 to 8). Labelling bins as distinct types of branches allowed for the estimation of up to eight different *d*_N_/*d*_S_ values per gene. Conjointly, for each binned model, an alternative null model assuming no difference in *d*_N_/*d*_S_ between branches was run (one-ratio model). The difference between models was compared using a likelihood-ratio test (LRT) by comparing twice the log-likelihood difference between the two models which is assumed to be *χ*^2^ distributed, with the relevant degrees of freedom (Yang 2007).

To estimate rate hetereogeneity among branches in noncoding regions, we used a model where we assumed equal rates among branches (e.g. a global clock, clock=1) and compared it to a model where we assumed different rates for the binned branches (clock=2), assessing significant differences between the models using a likelihood ratio test. For the simulations (Figure S2) we randomly chose a 222bp long genomic region with 67 species. We run a free branch model (clock=0) and used the obtained parameters as input for INDELible (Fletcher and Yang 2009). We simulated 100 sets of sequences and applied two types of binning: (1) A binning that grouped similar branch lengths and (2) an arbitrary binning. We considered 5 different numbers of bins (with 2,4,8,16 and unrestricted number of bins). We then conducted rate estimation on each of the binning approaches and calculated how well these estimates correlated (Kendall’s *τ* correlation coefficient) with the input parameters for INDELible (e.g. the simulation input) as well as the estimated values from the free branch model.

### Hotspot approach for the detection of genes under positive selection

To formally test for rapid and potentially positive selection on branches with increased rates of morphological evolution, we used a ‘hotspot’ approach. As opposed to focusing on large-effect genes important across distantly-related avian taxa, we identified and marked specific, individual branches undergoing the fastest rates of morphological evolution, according to rate estimates from Cooney et al. (2017). At these branches, we hypothesize to detect higher *d*_N_/*d*_S_ estimates relative to background branches.

### Phenotype and pathway ontologies, protein databases and statistical analyses

To determine the putative function of genes detected and enriched according to pathway and phenotype enrichment, we used WebGestalt (Wang et al. 2017) based on the human annotation. Specifically, we used the latest release of WebGestalt (last accessed 11.3.2019), and ran an Overrepresentation Enrichment Analysis (ORA) for phenotypes (Human Phenotype Ontology), pathways (Wikipathways) and diseases (Glad4U). We set the minimum number of genes for a category to 40 and reported top statistical significant results as weighted cover set (as implemented in WebGestalt). We also obtained a set of 511 genes known from mouse knock-out phenotypes to result in abnormal craniofacial morphology or development (Brunskill et al. 2014). To account for multiple testing in our binned and hotspot models, *χ*^2^-squared P-values were corrected using the Benjamini-Hochberg procedure (Benjamini and Hochberg 1995). We used Kendall’s *τ* correlation coefficient to compare the association between increasing bin number and corresponding *d*_N_/*d*_S_ (coding) and substitution rates (noncoding) for each gene. Statistical analysis was conducted using the SciPy library in Python, and graphs were produced using the ‘tidyverse’ package in R (Wilkinson 2011; R Core Team 2018) and the ‘matplotlib’ package in Python. Phylogenies were produced using the ‘phytools’ package in R (Revell 2012). Protein interaction partners for ALX1, BMP1 and CALM1 were retrieved from the STRING database (Szklarczyk et al. 2015) based on the human annotation requiring a minimum confidence score of 0.6 for all interaction partners. Motif detection was conducted using DREME (Bailey 2011) along with the identification of potential binding proteins using TOMTOM (Gupta et al. 2007). Specifically, we focused on vertebrate binding proteins using a common set of three available databases (JASPAR2018_CORE_vertebrates_non-redundant, jolma2013, uniprobe_mouse) that together contained 649 annotated motif binding proteins.

### Population genetic analysis in Darwin’s finches population with diverse beak morphology

We obtained per site measurements of population differentiation (fixation index *F*_ST_ and nucleotide diversity *θ* (Watterson 1975; Weir and Cockerham 1984) by calculating and contrasting genetic diversity of Darwin finch populations (Lamichhaney et al. 2015) with blunt (5 and 10 individuals from *Geospiza magnirostris* and *G. conirostris* populations, respectively) and pointed beaks (10 and 8 individuals from *G. conirostris* and *G. difficilis* populations, respectively).

### Software availability

Scripts concerning the analysis of protein coding regions and noncoding regions are available at GitHub (https://github.com/LeebanY/avian-comparative-genomics; https://githubcom/mattheatley/bird_project) and as Supplemental Code.

## Acknowledgements

We thank Sangeet Lamichhaney for providing *F_ST_* data on the Darwin finch populations and Victor Soria-Carrasco, Gavin Thomas and Jon Slate for constructive comments at earlier stages of this work. We also want to thank Ahmet Denli and three anonymous reviewers for their comments that helped to improve the quality of this work.

## Author Contributions

LY drafted the manuscript; TIG and CRC designed and supervised the study; TIG, LY, MH and JP conducted research; HJB contributed to research; TIG finalized the manuscript with written input from all authors

## References

Abzhanov A, Kuo WP, Hartmann C, Grant BR, Grant PR, Tabin CJ. 2006. The calmodulin pathway and evolution of elongated beak morphology in Darwin’s finches. Nature 442: 563–567.

Abzhanov A, Protas M, Grant BR, Grant PR, Tabin CJ. 2004. Bmp4 and morphological variation of beaks in Darwin’s finches. Science 305: 1462–1465. http://www.ncbi.nlm.nih.gov/pubmed/15353802.

Afanasyeva A, Bockwoldt M, Cooney CR, Heiland I, Gossmann TI. 2018. Human long intrinsically disordered protein regions are frequent targets of positive selection. Genome research 28: 975–982. http://www.ncbi.nlm.nih.gov/pubmed/29858274 http://www.pubmedcentral.nih.gov/articlerender.fcgi?artid=PMC6028134.

Andersson R, Sandelin A. 2020. Determinants of enhancer and promoter activities of regulatory elements. 21: 71–87.

Bailey TL. 2011. DREME: motif discovery in transcription factor ChIP-seq data. Bioinformatics 27: 1653–1659. https://academic.oup.com/bioinformatics/article-lookup/doi/10.1093/bioinformatics/btr261.

Balanoff AM, Bever GS, Rowe TB, Norell MA. 2013. Evolutionary origins of the avian brain. Nature 501: 93–96. http://dx.doi.org/10.1038/nature12424.

Benjamini Y, Hochberg Y. 1995. Controlling the false discovery rate: a practical and powerful approach to multiple testing. Journal of the Royal Statistical Society Series B (Methodological).

Berv JS, Field DJ. 2018. Genomic Signature of an Avian Lilliput Effect across the K-Pg Extinction. Systematic Biology 67: 1–13.

Bhullar AB-aS, Morris ZS, Sefton EM, Bhullar B-aS, Morris ZS, Sefton EM, Tok A, Tokita M, Namkoong B, Camacho J, et al. 2015. A molecular mechanism for the origin of a key evolutionary innovation, the bird beak and palate, revealed by an integrative approach to major transitions in vertebrate history. 1665–1677.

Bhullar B-AS, Hanson M, Fabbri M, Pritchard A, Bever GS, Hoffman E. 2016. How to Make a Bird Skull: Major Transitions in the Evolution of the Avian Cranium, Paedomorphosis, and the Beak as a Surrogate Hand. Integrative and Comparative Biology 56: 389–403. https://academic.oup.com/icb/article-lookup/doi/10.1093/icb/icw069.

Bhullar B-AS, Marugán-Lobón J, Racimo F, Bever GS, Rowe TB, Norell MA, Abzhanov A. 2012. Birds have paedomorphic dinosaur skulls. Nature 487: 223–226. http://www.nature.com/articles/nature11146.

Bielawski JP, Baker JL, Mingrone J. 2016. Inference of Episodic Changes in Natural Selection Acting on Protein Coding Sequences via CODEML. In Current protocols in bioinformatics, Vol. 54 of, pp. 6.15.1–6.15.32, John Wiley & Sons, Inc., Hoboken, NJ, USA http://doi.wiley.com/10.1002/cpbi.2.

Blanchette M, Kent WJ, Riemer C, Elnitski L, Smit AFA, Roskin KM, Baertsch R, Rosenbloom K, Clawson H, Green ED, et al. 2004. Aligning multiple genomic sequences with the threaded blockset aligner. Genome research 14: 708–15. http://www.ncbi.nlm.nih.gov/pubmed/15060014 http://www.pubmedcentral.nih.gov/articlerender.fcgi?artid=PMC383317.

Bolívar P, Guéguen L, Duret L, Ellegren H, Mugal CF. 2019. GC-biased gene conversion conceals the prediction of the nearly neutral theory in avian genomes. Genome Biology 20: 5. https://genomebiology.biomedcentral.com/articles/10.1186/s13059-018-1613-z.

Boĺivar P, Mugal CF, Nater A, Ellegren H. 2016. Recombination rate variation modulates gene sequence evolution mainly via GC-Biased gene conversion, not Hill-Robertson interference, in an avian system. Molecular Biology and Evolution 33: 216–227.

Bosse M, Spurgin LG, Laine VN, Cole EF, Firth JA, Gienapp P, Gosler AG, McMahon K, Poissant J, Verhagen I, et al. 2017a. Recent natural selection causes adaptive evolution of an avian polygenic trait. Science 358: 365–368.

Bosse M, Spurgin LG, Laine VN, Cole EF, Firth JA, Gienapp P, Gosler AG, McMahon K, Poissant J, Verhagen I, et al. 2017b. Recent natural selection causes adaptive evolution of an avian polygenic trait. Science.

Botero-Castro F, Figuet E, Tilak M-k, Nabholz B, Galtier N. 2017. Avian genomes revisited: hidden genes uncovered and the rates vs. traits paradox in birds. Molecular Biology and Evolution 1–9. http://academic.oup.com/mbe/article/doi/10.1093/molbev/msx236/4104406/Avian-genomes-revisited-hidden-genes-uncovered-and.

Boyle EA, Li YI, Pritchard JK. 2017. An Expanded View of Complex Traits: From Polygenic to Omnigenic. Cell 169: 1177–1186. https://www.sciencedirect.com/science/article/pii/S0092867417306293?via{\%}3Dihub.

Brugmann SA, Powder KE, Young NM, Goodnough LH, Hahn SM, James AW, Helms JA, Lovett M. 2010. Comparative gene expression analysis of avian embryonic facial structures reveals new candidates for human craniofacial disorders. Human molecular genetics 19: 920–30. http://www.ncbi.nlm.nih.gov/pubmed/20015954 http://www.pubmedcentral.nih.gov/articlerender.fcgi?artid=PMC2816616.

Brunskill EW, Potter AS, Distasio A, Dexheimer P, Plassard A, Aronow BJ, Potter SS. 2014. A gene expression atlas of early craniofacial development. Developmental biology 391: 133–46.

Butler JE, Kadonaga JT. 2002. The RNA polymerase II core promoter: A key component in the regulation of gene expression. 16: 2583–2592.

Capella-Gutierrez S, Silla-Martinez JM, Gabaldon T. 2009. trimAl: a tool for automated alignment trimming in large-scale phylogenetic analyses. Bioinformatics 25: 1972–1973. https://academic.oup.com/bioinformatics/article-lookup/doi/10.1093/bioinformatics/btp348.

Cooney CR, Bright JA, Capp EJR, Chira AM, Hughes EC, Moody CJA, Nouri LO, Varley ZK, Thomas GH. 2017. Mega-evolutionary dynamics of the adaptive radiation of birds. Nature 542: 344–347. http://www.nature.com/doifinder/10.1038/nature21074.

Corcoran P, Gossmann TI, Barton HJ, Consortium TGTH, Slate J, Zeng K. 2017. Determinants of the efficacy of natural selection on coding and noncoding variability in two passerine species. Genome Biology and Evolution 10: 1062–1062. http://academic.oup.com/gbe/article/9/11/2958/4583627 https://academic.oup.com/gbe/article/10/4/1062/4963733.

Ducy P, Desbois C, Boyce B, Pinero G, Story B, Dunstan C, Smith E, Bonadio J, Goldstein S, Gundberg C, et al. 1996. Increased bone formation in osteocalcin-deficient mice. 382: 448–452.

Felice RN, Goswami A. 2017. Developmental origins of mosaic evolution in the avian cranium. Proceedings of the National Academy of Sciences 115: 201716437. http://www.pnas.org/lookup/doi/10.1073/pnas.1716437115.

Figuet E, Bonneau M, Carrio EM, Nadachowska-brzyska K, Ellegren H, Galtier N. 2017. Life History Traits, Protein Evolution, and the Nearly Neutral Theory in Amniotes. 33: 1517–1527.

Fisher R a. 1930. The Genetical Theory of Natural Selection. Genetics 154: 272. http://openlibrary.org/books/OL7084333M.

Fletcher W, Yang Z. 2009. INDELible: A Flexible Simulator of Biological Sequence Evolution. Molecular Biology and Evolution 26: 1879–1888. https://academic.oup.com/mbe/article-lookup/doi/10.1093/molbev/msp098.

Foote M. 1997. THE EVOLUTION OF MORPHOLOGICAL DIVERSITY. Annual Review of Ecology and Systematics 28: 129–152. http://linkinghub.elsevier.com/retrieve/pii/0142694X80900496 http://www.annualreviews.org/doi/10.1146/annurev.ecolsys.28.1.129.

Galtier N, Duret L, Glémin S, Ranwez V. 2009. GC-biased gene conversion promotes the fixation of deleterious amino acid changes in primates. 25: 1–5.

Gossmann TI, Bockwoldt M, Diringer L, Schwarz F, Schumann V-F. 2018. Evidence for Strong Fixation Bias at 4-fold Degenerate Sites Across Genes in the Great Tit Genome. Frontiers in Ecology and Evolution 6: 203. https://www.frontiersin.org/article/10.3389/fevo.2018.00203/full.

Gossmann T, Song B-H, Windsor A, Mitchell-Olds T, Dixon C, Kapralov M, Filatov D, Eyre-Walker A. 2010. Genome wide analyses reveal little evidence for adaptive evolution in many plant species. Molecular Biology and Evolution 27.

Gossmann T, Waxman D, Eyre-Walker A. 2014. Fluctuating selection models and Mcdonald-Kreitman type analyses. PLoS ONE 9.

Grant BR, Grant PR. 1996. High Survival of Darwin’s Finch Hybrids: Effects of Beak Morphology and Diets. Ecological Society of America 77: 500–509. http://www.jstor.org/stable/2265625.

Grant PR, Grant BR. 2009. The secondary contact phase of allopatric speciation in Darwin’s finches. Proceedings of the National Academy of Sciences 106: 20141–20148. http://www.pnas.org/cgi/doi/10.1073/pnas.0911761106.

Gupta S, Stamatoyannopoulos JA, Bailey TL, Noble W. 2007. Quantifying similarity between motifs. Genome Biology 8: R24. http://genomebiology.biomedcentral.com/articles/10.1186/gb-2007-8-2-r24.

Hackett SJ, Kimball RT, Reddy S, Bowie RCK, Braun EL, Braun MJ, Chojnowski JL, Cox WA, Han K-L, Harshman J, et al. 2008. A Phylogenomic Study of Birds Reveals Their Evolutionary History. Science 320: 1763–1768. http://www.sciencemag.org/cgi/doi/10.1126/science.1157704.

Han F, Lamichhaney S, Grant BR, Grant PR, Andersson L, Webster MT. 2017. Gene flow, ancient polymorphism, and ecological adaptation shape the genomic landscape of divergence among Darwin’s finches. 1–12.

Hill WG. 2010. Understanding and using quantitative genetic variation. Philosophical Transactions of the Royal Society B: Biological Sciences 365: 73–85.

Hu Z, Sackton TB, Edwards SV, Liu JS. 2019. Bayesian Detection of Convergent Rate Changes of Conserved Noncoding Elements on Phylogenetic Trees ed. S.K. Pond. Molecular Biology and Evolution 36: 1086–1100. https://academic.oup.com/mbe/article/36/5/1086/5372678.

Huber SK, Podos J. 2006. Beak morphology and song features covary in a population of Darwin’s finches (Geospiza fortis). Biological Journal of the Linnean Society 88: 489–498.

Huerta-Sanchez E, Durrett R, Bustamante CD. 2008. Population genetics of polymorphism and divergence under fluctuating selection. Genetics 178: 325–37. http://www.ncbi.nlm.nih.gov/pubmed/17947441 http://www.pubmedcentral.nih.gov/articlerender.fcgi?artid=PMC2206081.

Hughes M, Gerber S, Wills MA. 2013. Clades reach highest morphological disparity early in their evolution. Proceedings of the National Academy of Sciences 110: 13875–13879. http://www.pnas.org/lookup/doi/10.1073/pnas.1302642110.

Hunter JP. 1998. Key innovations and the ecology of macroevolution. Trends in ecology & evolution 13: 31–6. http://www.ncbi.nlm.nih.gov/pubmed/21238187.

Jones CT, Youssef N, Susko E, Bielawski JP. 2016. Shifting Balance on a Static Mutation–Selection Landscape: A Novel Scenario of Positive Selection. Molecular Biology and Evolution 34: msw237. https://academic.oup.com/mbe/article-lookup/doi/10.1093/molbev/msw237.

Karsenty G, Ferron M. 2012. The contribution of bone to whole-organism physiology. Nature 481: 314–320.

Kimura M. 1983. The Neutral Theory of Molecular Evolution. http://books.google.co.il/books?id=olIoSumPevYC{\&}lpg=PP1{\&}dq=theneutraltheoryofmolecularevolution{\%}5Cnhttps://books.google.com/books?hl=en{\&}lr={\&}id=olIoSumPevYC{\&}pgis=1.

Lahti DC, Johnson NA, Ajie BC, Otto SP, Hendry AP, Blumstein DT, Coss RG, Donohue K, Foster SA. 2009. Relaxed selection in the wild. Trends in Ecology and Evolution 24: 487–496.

Lamichhaney S, Berglund J, Almén MS, Maqbool K, Grabherr M, Martinez-Barrio A, Promerová M, Rubin C-J, Wang C, Zamani N, et al. 2015. Evolution of Darwin’s finches and their beaks revealed by genome sequencing. Nature 518: 371–375. http://www.nature.com/doifinder/10.1038/nature14181.

Lamichhaney S, Card DC, Grayson P, Tonini JFR, Bravo GA, Näpflin K, Termignoni-Garcia F, Torres C, Burbrink F, Clarke JA, et al. 2019. Integrating natural history collections and comparative genomics to study the genetic architecture of convergent evolution. Philosophical Transactions of the Royal Society B: Biological Sciences 374: 20180248. https://royalsocietypublishing.org/doi/10.1098/rstb.2018.0248.

Lamichhaney S, Han F, Webster MT, Andersson L, Grant BR, Grant PR. 2018. Rapid hybrid speciation in Darwin’s finches. Science 359: 224–228. http://www.sciencemag.org/lookup/doi/10.1126/science.aao4593.

Lanfear R, Ho SYW, Love D, Bromham L. 2010. Mutation rate is linked to diversification in birds. 107: 20423–20428.

Lanfear R, Kokko H, Eyre-walker A. 2014. Population size and the rate of evolution. Trends in Ecology & Evolution 29: 33–41. http://dx.doi.org/10.1016/j.tree.2013.09.009.

Lartillot N. 2013. Interaction between selection and biased gene conversion in mammalian protein-coding sequence evolution revealed by a phylogenetic covariance analysis. Molecular Biology and Evolution 30: 356–368.

Lee NK, Sowa H, Hinoi E, Ferron M, Ahn JD, Confavreux C, Dacquin R, Mee PJ, McKee MD, Jung DY, et al. 2007. Endocrine Regulation of Energy Metabolism by the Skeleton. Cell 130: 456–469.

Lefebvre V, Li P, De Crombrugghe B. 1998. A new long form of Sox5 (L-Sox5), Sox6 and Sox9 are coexpressed in chondrogenesis and cooperatively activate the type II collagen gene. EMBO Journal 17: 5718–5733.

Levy Karin E, Wicke S, Pupko T, Mayrose I. 2017. An Integrated Model of Phenotypic Trait Changes and Site-Specific Sequence Evolution. Systematic Biology 66: 917–933. http://academic.oup.com/sysbio/article/66/6/917/2978030/An-Integrated-Model-of-Phenotypic-Trait-Changes.

Li L, Stoeckert CJJ, Roos DS. 2003. OrthoMCL: Identification of Ortholog Groups for Eukaryotic Genomes. Genome Research 13: 2178–2189. http://genome.cshlp.org/cgi/content/full/13/9/2178.

Lloyd SP. 1982. Least Squares Quantization in PCM. IEEE Transactions on Information Theory.

Lowe CB, Clarke JA, Baker AJ, Haussler D, Edwards SV. 2015. Feather Development Genes and Associated Regulatory Innovation Predate the Origin of Dinosauria. Molecular Biology and Evolution 32: 23–28. https://academic.oup.com/mbe/article-lookup/doi/10.1093/molbev/msu309.

Löytynoja A, Goldman N. 2008. Phylogeny-aware gap placement prevents errors in sequence alignment and evolutionary analysis. Science 320: 1632–1635.

Lynch M. 1987. The consequences of fluctuating selection for isozyme polymorphisms in Daphnia. Genetics 115: 657–669.

Machado JP, Johnson WE, Gilbert MTP, Zhang G, Jarvis ED, O’Brien SJ, Antunes A. 2016. Bone-associated gene evolution and the origin of flight in birds. BMC Genomics 17: 1–15. http://dx.doi.org/10.1186/s12864-016-2681-7.

Mallarino R, Grant PR, Grant BR, Herrel A, Kuo WP, Abzhanov A. 2011. Two developmental modules establish 3D beak-shape variation in Darwin’s finches. Proceedings of the National Academy of Sciences 108: 4057–4062. http://www.pnas.org/cgi/doi/10.1073/pnas.1011480108.

Manceau M, Domingues VS, Linnen CR, Rosenblum EB, Hoekstra HE. 2010. Convergence in pigmentation at multiple levels: mutations, genes and function. Philosophical Transactions of the Royal Society B: Biological Sciences 365: 2439–2450. https://royalsocietypublishing.org/doi/10.1098/rstb.2010.0104.

Matsumoto T, Akashi H, Yang Z. 2015. Evaluation of Ancestral Sequence Reconstruction Methods to Infer Nonstationary Patterns of Nucleotide Substitution. Genetics 200: 873–90. http://www.genetics.org/lookup/doi/10.1534/genetics.115.177386.

Mayrose I, Otto SP. 2011. A Likelihood Method for Detecting Trait-Dependent Shifts in the Rate of Molecular Evolution. Molecular Biology and Evolution 28: 759–770. https://academic.oup.com/mbe/article-lookup/doi/10.1093/molbev/msq263.

Mcentee JP, Tobias JA, Sheard C, Burleigh JG. 2018. Tempo and timing of ecological trait divergence associated with transitions to coexistence in birds. Nature Ecology & Evolution 2: 1–17. http://www.biorxiv.org/content/biorxiv/early/2017/04/26/083253.full.pdf.

Mera P, Laue K, Ferron M, Confavreux C, Wei J, Galán-Díez M, Lacampagne A, Mitchell SJ, Mattison JA, Chen Y, et al. 2016. Osteocalcin Signaling in Myofibers Is Necessary and Sufficient for Optimum Adaptation to Exercise. Cell Metabolism 23: 1078–1092. http://linkinghub.elsevier.com/retrieve/pii/S1550413116302224.

Merkin J, Russell C, Chen P, Burge CB. 2012. Evolutionary Dynamics of Gene and Isoform Regulation in Mammalian Tissues. Science 338: 1593–1599. http://www.sciencemag.org/cgi/doi/10.1126/science.1228186.

Merrill AE, Eames BF, Weston SJ, Heath T, Schneider RA. 2008. Mesenchyme-dependent BMP signaling directs the timing of mandibular osteogenesis. Development 135: 1223–1234. http://www.ncbi.nlm.nih.gov/pubmed/18287200 http://www.pubmedcentral.nih.gov/articlerender.fcgi?artid=PMC2844338 http://dev.biologists.org/cgi/doi/10.1242/dev.015933.

Nabholz B, Lanfear R, Fuchs J. 2016. Body mass-corrected molecular rate for bird mitochondrial DNA. Molecular Ecology 25: 4438–4449.

Nabholz B, Uwimana N, Lartillot N. 2013. Reconstructing the phylogenetic history of long-term effective population size and life-history traits using patterns of amino acid replacement in mitochondrial genomes of mammals and birds. Genome Biology and Evolution 5: 1273–1290.

O’Connor TD, Mundy NI. 2013. Evolutionary Modeling of Genotype-Phenotype Associations, and Application to primate coding and Non-coding mtDNA Rate Variation. Evolutionary Bioinformatics 9: EBO.S11600. http://journals.sagepub.com/doi/10.4137/EBO.S11600.

O’Connor TD, Mundy NI. 2009. Genotype–phenotype associations: substitution models to detect evolutionary associations between phenotypic variables and genotypic evolutionary rate. Bioinformatics 25: i94–i100. https://academic.oup.com/bioinformatics/article-lookup/doi/10.1093/bioinformatics/btp231.

O’Hara RB. 2005. Comparing the effects of genetic drift and fluctuating selection on genotype frequency changes in the scarlet tiger moth. Proceedings of the Royal Society B: Biological Sciences 272: 211–217.

Okita K, Yamanaka S. 2006. Intracellular signaling pathways regulating pluripotency of embryonic stem cells. Current stem cell research & therapy 1: 103–11. http://www.ncbi.nlm.nih.gov/pubmed/18220859.

O’Leary NA, Wright MW, Brister JR, Ciufo S, Haddad D, McVeigh R, Rajput B, Robbertse B, Smith-White B, Ako-Adjei D, et al. 2016. Reference sequence (RefSeq) database at NCBI: current status, taxonomic expansion, and functional annotation. Nucleic Acids Research 44: D733–D745. https://academic.oup.com/nar/article-lookup/doi/10.1093/nar/gkv1189.

Östlund G, Schmitt T, Forslund K, Köstler T, Messina DN, Roopra S, Frings O, Sonnhammer EL. 2009. Inparanoid 7: New algorithms and tools for eukaryotic orthology analysis. Nucleic Acids Research 38: 196–203.

Parsons KJ, Albertson RC. 2009. Roles for Bmp4 and CaM1 in Shaping the Jaw: Evo-Devo and Beyond. Annual Review of Genetics 43: 369–388. http://www.annualreviews.org/doi/10.1146/annurev-genet-102808-114917.

Piechota M, Korostynski M, Przewlocki R. 2010. Identification of cis-Regulatory Elements in the Mammalian Genome: The cREMaG Database ed. C.A. Ouzounis. PLoS ONE 5: e12465. https://dx.plos.org/10.1371/journal.pone.0012465.

Podos J. 2001. Correlated evolution of morphology and vocal structure in Darwin’s finches. Nature 409: 185–188.

Prudent X, Parra G, Schwede P, Roscito JG, Hiller M. 2016. Controlling for Phylogenetic Relatedness and Evolutionary Rates Improves the Discovery of Associations Between Species’ Phenotypic and Genomic Differences. Molecular Biology and Evolution 33: 2135–2150. https://academic.oup.com/mbe/article-lookup/doi/10.1093/molbev/msw098.

Prum RO, Berv JS, Dornburg A, Field DJ, Townsend JP, Lemmon EM, Lemmon AR. 2015. A comprehensive phylogeny of birds (Aves) using targeted next-generation DNA sequencing. Nature 526: 569–573. http://www.nature.com/doifinder/10.1038/nature15697.

Raymond MH, Schutte BC, Torner JC, Burns TL WM. 1999. Osteocalcin: genetic and physical mapping of the human gene BGLAP and its potential role in postmenopausal osteoporosis. Genomics 60: 210–17.

R Core Team. 2018. R: A Language and Environment for Statistical Computing. R Foundation for Statistical Computing, Vienna, Austria https://www.r-project.org/.

Revell LJ. 2012. phytools: An R package for phylogenetic comparative biology (and other things). Methods in Ecology and Evolution.

Ricklefs RE. 2004. Cladogenesis and morphological diversification in passerine birds. Nature.

Rockman MV. 2012. The QTN program and the alleles that matter for evolution: All that’s gold does not glitter. Evolution 66: 1–17.

Rosenblum EB, Parent CE, Brandt EE. 2014. The Molecular Basis of Phenotypic Convergence. Annual Review of Ecology, Evolution, and Systematics 45: 203–226. http://www.annualreviews.org/doi/10.1146/annurev-ecolsys-120213-091851.

Sackton TB, Grayson P, Cloutier A, Hu Z, Liu JS, Wheeler NE, Gardner PP, Clarke JA, Baker AJ, Clamp M, et al. 2019. Convergent regulatory evolution and loss of flight in paleognathous birds. Science (New York, NY) 364: 74–78. http://www.ncbi.nlm.nih.gov/pubmed/30948549.

Schluter D. 2000. The Ecology of Adaptive Radiation.

Seki R, Li C, Fang Q, Hayashi S, Egawa S, Hu J, Xu L, Pan H, Kondo M, Sato T, et al. 2017. Functional roles of Aves class-specific cis-regulatory elements on macroevolution of bird-specific features. Nature Communications 8: 14229. http://www.nature.com/doifinder/10.1038/ncomms14229.

Sharma V, Hecker N, Roscito JG, Foerster L, Langer BE, Hiller M. 2018. A genomics approach reveals insights into the importance of gene losses for mammalian adaptations. Nature Communications 9: 1215. http://www.nature.com/articles/s41467-018-03667-1.

Shultz AJ, Sackton TB. 2019. Immune genes are hotspots of shared positive selection across birds and mammals. eLife 8.

Simpson GG. 1944. Tempo and mode in evolution. Columbia Univ. Press, New York https://www.worldcat.org/title/tempo-and-mode-in-evolution/oclc/993515.

Stern DL. 2013. The genetic causes of convergent evolution. Nature Reviews Genetics 14: 751–764. http://www.nature.com/articles/nrg3483.

Stroud JT, Losos JB. 2016. Ecological Opportunity and Adaptive Radiation. Annual Review of Ecology, Evolution, and Systematics 47: 507–532. http://www.annualreviews.org/doi/10.1146/annurev-ecolsys-121415-032254.

Szklarczyk D, Franceschini A, Wyder S, Forslund K, Heller D, Huerta-Cepas J, Simonovic M, Roth A, Santos A, Tsafou KP, et al. 2015. STRING v10: Protein-protein interaction networks, integrated over the tree of life. Nucleic Acids Research 43: D447–D452.

Venditti C, Meade A, Pagel M. 2011. Multiple routes to mammalian diversity. Nature 479: 393–396. http://dx.doi.org/10.1038/nature10516.

Wang J, Vasaikar S, Shi Z, Greer M, Zhang B. 2017. WebGestalt 2017: a more comprehensive, powerful, flexible and interactive gene set enrichment analysis toolkit. Nucleic Acids Research 45: W130–W137. https://academic.oup.com/nar/article-lookup/doi/10.1093/nar/gkx356.

Watterson G. 1975. On the number of segregating sites in genetical models without recombination. Theoretical Population Biology 7: 256–276. https://www.sciencedirect.com/science/article/pii/0040580975900209?via{\%}3Dihub.

Weber CC, Nabholz B, Romiguier J, Ellegren H. 2014. Kr/Kc but not dN/dS correlates positively with body mass in birds, raising implications for inferring lineage-specific selection. 1–13.

Weir BS, Cockerham CC. 1984. Estimating F-Statistics for the Analysis of Population Structure. Evolution 38: 1358. https://www.jstor.org/stable/2408641?origin=crossref.

Whitney O, Pfenning AR, Howard JT, Blatti CA, Liu F, Ward JM, Wang R, Audet J-N, Kellis M, Mukherjee S, et al. 2014. Core and region-enriched networks of behaviorally regulated genes and the singing genome. Science 346: 1256780–1256780. http://www.sciencemag.org/cgi/doi/10.1126/science.1256780.

Wilkinson L. 2011. ggplot2: Elegant Graphics for Data Analysis by WICKHAM, H. Biometrics.

Wirthlin M, Lovell PV, Jarvis ED, Mello CV. 2014. Comparative genomics reveals molecular features unique to the songbird lineage. BMC Genomics 15: 1082. http://bmcgenomics.biomedcentral.com/articles/10.1186/1471-2164-15-1082.

Wittkopp PJ, Kalay G. 2012. Cis-regulatory elements: Molecular mechanisms and evolutionary processes underlying divergence. 13: 59–69.

Wright D, Boije H, Meadows JRS, Bed’hom B, Gourichon D, Vieaud A, Tixier-Boichard M, Rubin C-J, Imsland F, Hallböök F, et al. 2009. Copy Number Variation in Intron 1 of SOX5 Causes the Pea-comb Phenotype in Chickens ed. D.L. Stern. PLoS Genetics 5: e1000512. http://dx.plos.org/10.1371/journal.pgen.1000512.

Wu P, Jiang T-X, Suksaweang S, Widelitz RB, Chuong C-M. 2004. Molecular shaping of the beak. Science (New York, NY) 305: 1465–6. http://www.ncbi.nlm.nih.gov/pubmed/15353803 http://www.pubmedcentral.nih.gov/articlerender.fcgi?artid=PMC4380220.

Xu X, Zhou Z, Dudley R, MacKem S, Chuong CM, Erickson GM, Varricchio DJ. 2014. An integrative approach to understanding bird origins. Science 346.

Yang Z. 1998. Likelihood ratio tests for detecting positive selection and application to primate lysozyme evolution. Molecular Biology and Evolution 15: 568–573. https://academic.oup.com/mbe/article-lookup/doi/10.1093/oxfordjournals.molbev.a025957.

Yang Z. 2007. PAML 4: Phylogenetic analysis by maximum likelihood. Molecular Biology and Evolution 24: 1586–1591.

Yang Z, Nielsen R. 1998. Synonymous and nonsynonymous rate variation in nuclear genes of mammals. Journal of Molecular Evolution 46: 409–418.

Yang Z, Nielsen R, Hasegawa M. 1998. Models of Amino Acid Substitution and Applications to Mitochondrial Protein Evolution. Mol Biol Evol 15: 1600–1611.

