## Supplemental Methods for "Noncoding regions underpin avian bill shape diversification at macroevolutionary scales"

### Supplemental Material

**Supplemental Methods** **1**

**Supplemental References** **16**

### Supplemental Methods

##### Multiple sequence alignments for protein coding genes

We used genomes of 57 bird species with high quality annotations from NCBI RefSeq (O'Leary et al. 2016) (Table S2). First, 12,013 orthologous protein coding genes were retrieved using RefSeq and HGNC gene identifiers, alongside reciprocal BLAST approaches based on three focal species, chicken, great tit and zebra finch - three of the best annotated high quality bird genomes available to date (Li et al. 2003; Östlund et al. 2009; Afanasyeva et al. 2018). We then performed a first set of alignment runs using PRANK (Löytynoja and Goldman 2008). To ensure the quality of these sequence alignments, we applied a customised pipeline. Firstly, alignments were filtered for length and the number of species they contained. Generally, we applied a length filter that removed alignments containing more than 1500 amino acid residues (for computational reasons) and less than 50 amino acid residues (for power reasons). These thresholds were based on distributions of overall sequence length across all alignments. Furthermore, we compared the number of gaps and length of sequences to a reference sequence in all alignments. Regarded as the most well-annotated, high-quality avian genome, we selected the red jungle fowl (i.e. chicken, *Gallus gallus*) as our reference sequence for all alignments (Hillier et al. 2004; Warren et al. 2017). Sequences determined to be too dissimilar (e.g. because of falsely aligning non-homologous regions within a protein), based on gappyness – the amount of gaps in a sequence – and overall sequence length, relative to our reference species sequence, were also removed from alignments. Specifically, for gappyness, if gaps resulted in more than 20% dissimilarity with our reference sequence, sequences were removed. We also limited the analyses to alignments containing 20 or more species, and removed alignments that did not contain the reference chicken sequence.

For reliable estimation of sequence divergence at the protein level, sequences that appeared too divergent were removed - caused by either elevated local mutation rate or, more likely, by falsely assigned orthologies. For this pairwise estimation of $d_{\mathrm{N}}$, $d_{\mathrm{S}}$ and $d_{\mathrm{N}}/d_{\mathrm{S}}$ was performed to determine saturation of non-synonymous and synonymous substitutions. Removal of saturated sequences was accomplished in two ways. First, pairwise synonymous substitution rates deemed too large were removed ($d_{\mathrm{S}}$>5), and second, if synonymous substitution rates exceeded twice the pairwise synonymous substitution rate between *G. gallus* and *Taeniopygia guttata* (zebra finch), two of our focal species, sequences were removed. In addition, if the pairwise non-synonymous substitution rate, $d_{\mathrm{N}}$, and the non-synonymous to synonymous substitution rate, $d_{\mathrm{N}}/d_{\mathrm{S}}$, exceeded two ($d_{\mathrm{N}}$>2 and $d_{\mathrm{N}}/d_{\mathrm{S}}$>2) sequences were removed from alignments. By this the sequences are conservatively aligned which reduces the chances of alignment-error signal of (positive) selection.

After a second alignment step with PRANK, to ensure positional homology, we utilised two masking programs: GBLOCKS and ZORRO. GBLOCKS calculates and uses positional homology to determine contiguous segments that are not conserved (Talavera and Castresana 2007). Additionally, GBLOCKS accounts for rapidly-evolving, homologous positions and flanking positional homology. For this, we used the following parameters to identify and remove unreliable positions: -t=p, -k=y, n=y, v =32000, -p=t. To supplement this, we used ZORRO, a probabilistic masking program which calculates posterior probabilities to determine the reliability of positions (Wu et al. 2012). Posterior probabilities calculated by ZORRO translate into scores that range from 0 to 10 – the higher the score, the better the positional homology. Positions that scored below 9 were removed from sequences. The removal of unreliable positions from sequences was performed with PAL2NAL (Suyama et al. 2006) using a customised script. Equally, PAL2NAL generated for each protein alignment the corresponding codon alignment in preparation for evolutionary analyses. A final length filter was applied to remove any alignments with a sequence length below 50 amino acids.

##### Rates of morphological beak shape evolution

Information on beak shape evolution was extracted from a recent study (Cooney et al. 2017) that quantified patterns of beak shape evolution across 2,028 species (>97% extant avian genera) covering the entire breadth of the avian clade. Briefly, this study used geometric morphometric data based on 3-D scans of museum specimens and multivariate rate heterogeneous models of trait evolution (Venditti et al. 2011) to estimate rates of beak shape evolution for all major branches in the avian phylogeny. Importantly, the beak shape measurements derived from this study are independent of variation in beak size, the effects of which are removed as part of standard geometric morphometric analyses (see Cooney et al. (2017) for full details). This is useful for our purposes as beak size tends to be strongly related to body size (which is known to covary with several genetic parameters), and because beak shape (rather than size) represents a key axis of ecomorphological differentiation between major avian groups (Cooney et al. 2017). To extract rate estimates for the species included in this study, we first pruned the 2,028 tip morphology rate-scaled phylogenies derived from Cooney et al. (2017) (based on the Hackett et al. (Hackett et al. 2008) backbone) to include only species for which coding/genomic information was available. We then divided the branch lengths in this pruned morphology rate-scaled tree by time (i.e. branch lengths from a similarly pruned time tree, also derived from Cooney et al. (2017)), to generate rate estimates specific to each branch in the pruned subtree. It is worth noting that our approach of pruning the

2,028 tip morphology rate-scaled tree is preferable to running a separate rates analysis including only a limited number of species included in our genomic dataset because the increased density of sampling in the larger tree will permit more accurate estimation of the magnitude and phylogenetic position of rate shifts in beak shape evolution across branches of the phylogeny.

##### Binned branch approach for the detection of large-effect genes and regulatory regions

To detect genes that may be undergoing repeated periods of rapid, possibly adaptive, evolution across multiple lineages, we grouped branches in each alignment phylogeny according to their rates of morphological evolution using *k*-means binning (Lloyd 1982). Here, we opted for up to eight (coding) and 16 (ASCHE) bins, respectively, to enable robust statistical analysis but still reasonable computational time for the substitution rate analysis. To phylogenetically link the genetic data to the morphological data we relied on the Hackett et al. backbone (Hackett et al. 2008), hence we did not account for phylogenetic heterogeneity among genes and possible gene-tree species tree discordance. Branches were grouped incrementally based on rates of trait evolution using a *k*-means binning approach, with the first bin representing branches with the slowest rates of morphological evolution, and the last bin representing branches with the fastest rates of morphological evolution (Figure 1). We assumed that genes involved in beak shape evolution would experience evolutionary rate change at the protein level ($d_{\mathrm{N}}/d_{\mathrm{S}}$) proportional to their respective rate of morphological evolution. Theoretically, we hypothesize that genes important in beak shape evolution across many branches would show a strong positive correlation.

In our analysis, we tested this using a branch model which assumes different substitution rates ($d_{\mathrm{N}}/d_{\mathrm{S}}$) across different, pre-defined, branches in a phylogeny using codeml (Yang 2007). Critically, the branch model may be useful in the detection of adaptive evolution occurring on particular branches (Yang et al. 1998; Yang 1998). Furthermore, we selected the branch model due to computational efficiency; the branch-site model and free-ratio model was deemed computationally intractable for a phylogeny of up to 57 species. Branches in each alignment's phylogeny were marked according to their respective bins (typically ranging from 1 to 8). Labelling bins as distinct types of branches allowed for the estimation of up to eight different $d_{\mathrm{N}}/d_{\mathrm{S}}$ values per gene. Conjointly, for each binned model, an alternative null model assuming no difference in $d_{\mathrm{N}}/d_{\mathrm{S}}$ between branches was run (one-ratio model). The difference between models was compared using a likelihood-ratio test (LRT) by comparing twice the log-likelihood difference between the two models which is assumed to be $\chi^2$ distributed, with the relevant degrees of freedom (Yang 2007) (e.g, seven degrees of freedom in case eight different branch categories were classified). If the binned model showed a significant difference to the one-ratio model an association between beak shape change and molecular rate change was inferred.

To estimate rate heterogeneity among branches in noncoding regions, we used a model where we assumed equal rates among branches (e.g. a global clock, clock=1) and compared it to a model where we assumed different rates for the binned branches (clock=2), assessing significant differences between the models using a likelihood ratio test using baseml from the paml package (Yang 2007). For the simulations (Figure S2) we randomly chose a 222bp long genomic region with 67 species. We run a free branch model (clock=0) and used the obtained parameters as input for INDELible (Fletcher and Yang 2009). We simulated 100 sets of sequences and applied two types of binning: (1) A binning that grouped similar branch lengths and (2) an arbitrary binning. We considered 5 different numbers of bins (with 2,4,8,16 and unrestricted number of bins). We then conducted rate estimation on each of the binning approaches and calculated how well these estimates correlated (Kendall's $\tau$ correlation coefficient) with the input parameters for INDELible (e.g. the simulation input) as well as the estimated values from the free branch model.

##### Hotspot approach for the detection of genes under positive selection

For each alignment we generated and conducted three independent branch models, corresponding to the three most rapidly evolving branches in each phylogeny. A null model assuming no differences in $d_{\mathrm{N}}/d_{\mathrm{S}}$ across branches in the phylogeny was conjointly computed. Again, the LRT was calculated to determine whether differences between each 'hotspot' model and the null model were significant. It is important to note that branches are not uniformly selected across alignments and alignment trees. This is because alignments vary in the number of species and branches they contain due to the filtering process applied. Hence, the selection of branches is dependent on species rates of morphological evolution relative to other species – the exclusion of species, particularly rapidly-evolving branches, causes new branches to be recruited in the hotspot-branch model. In total, five different branches rotate over our three hotspots (Figure S3). This can be done because each analysis is conducted per gene on the correctly pruned phylogeny. In most cases our fastest branch was an internal branch leading to the diversification of swifts (Apodidae), nightjars and their allies, (Caprimulgidae) and hummingbirds (Trochilidae). This is plausible given the disparity in beak shape, physiology and ecology that has arisen in this clade (Prum et al. 2015; Cooney et al. 2017).

##### Phenotype and pathway ontologies, protein databases and statistical analyses

To determine the putative function of genes detected and enriched according to pathway and phenotype enrichment, we used WebGestalt (Wang et al. 2017) based on the human annotation. Specifically, we used the latest release of WebGestalt (last accessed 11.3.2019), and ran an Overrepresentation Enrichment Analysis (ORA) for phenotypes (Human Phenotype Ontology), pathways (Wikipathways) and diseases (Glad4U). We set the minimum number of genes for a category to 40 and reported top statistical significant results as weighted cover set (as implemented in WebGestalt). We also obtained a set of 511 genes known from mouse knock-out phenotypes to result in abnormal craniofacial morphology or development (Brunskill et al. 2014). To account for multiple testing in our binned and hotspot models, $\chi^2$-squared P-values were corrected using the Benjamini-Hochberg procedure (Benjamini and Hochberg 1995). We used Kendall's $\tau$ correlation coefficient to compare the association between increasing bin number and corresponding $d_{\mathrm{N}}/d_{\mathrm{S}}$ (coding) and substitution rates (noncoding) for each gene. Statistical analysis was conducted using the SciPy library in Python, and graphs were produced using the 'tidyverse' package in R (Wilkinson 2011; R Core Team 2018) and the 'matplotlib' package in Python. Phylogenies were produced using the 'phytools' package in R (Revell 2012). Protein interaction partners for ALX1, BMP1 and CALM1 were retrieved from the STRING database (Szklarczyk et al. 2015) based on the human annotation requiring a minimum confidence score of 0.6 for all interaction partners. Motif detection was conducted using DREME (Bailey 2011) along with the identification of potential binding proteins using TOMTOM (Gupta et al. 2007). Specifically, we focused on vertebrate binding proteins using a common set of three available databases (JASPAR2018_CORE_vertebrates_non-redundant, jolma2013, uniprobe_mouse) that together contained 649 annotated motif binding proteins.

**Population genetic analysis in Darwin's finches population with diverse beak morphology**

We used the 39,806 noncoding genomic loci as focal regions and 1000 bp on either site of their center. To map our identified genomic loci onto the medium ground finch (*Geospiza fortis*) reference genome (Zhang et al. 2014), we used the best BLAST (default parameters) hit per region. We also extracted the same number of size and chromosome matched genomic regions that did not show an association with beak shape morphological diversification as control regions. To study the effect of selection at the focal and nearby sites due to linkage, a sliding window approach was used, applying a window size of 400bp every 50 bp around the center of the focal regions (Other window and step sizes gave very similar results). For $F_{\mathrm{ST}}$ we used the highest per site $F_{\mathrm{ST}}$ value for a particular genomic region in a given window and calculated the mean across all regions. Watterson's $\theta$ was calculated per genomic region in a given window and then averaged across all loci.

### Supplemental Figures

#### Supplemental Figure S1

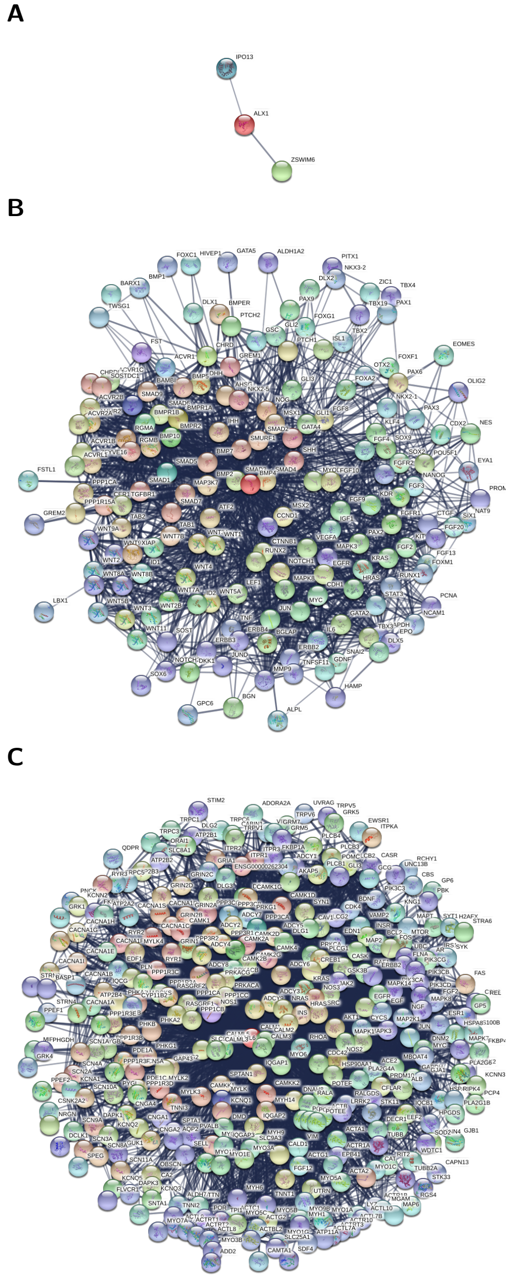

Figure S1: ***In-silico* interaction networks derived from the STRING database** for three proteins previously shown to be involed in the development of beak shape morphology.

#### Supplemental Figure S2

**A**

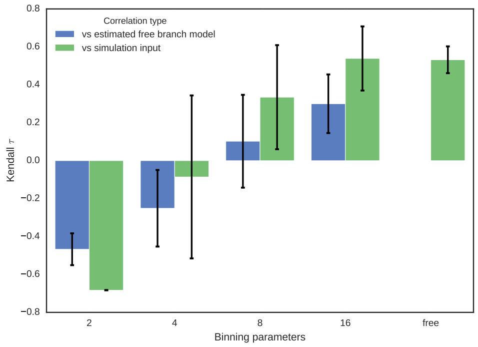

**B**

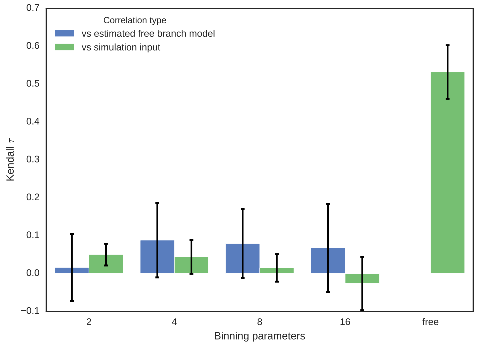

**C**

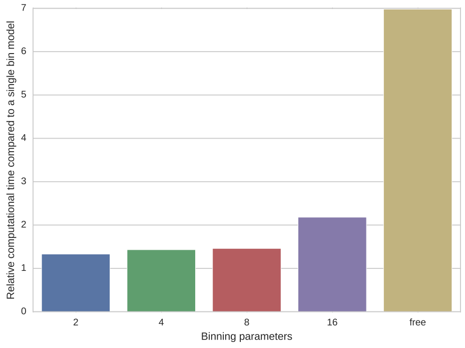

Figure S2: **Simulations to capture rate hetereogenity among branches by co-estimating rates of molecular change for grouped branches, estimated for noncoding regions.** (A) Correlation coefficients (Kendall $\tau$) of simulated and estimated rate hetereogeneity for different bin numbers, where branches of similar rates are grouped together. (B) Same approach using an arbitrary binning of branches (C) Relative computational time requirements for different number of bins.

**Supplemental Figure S3**

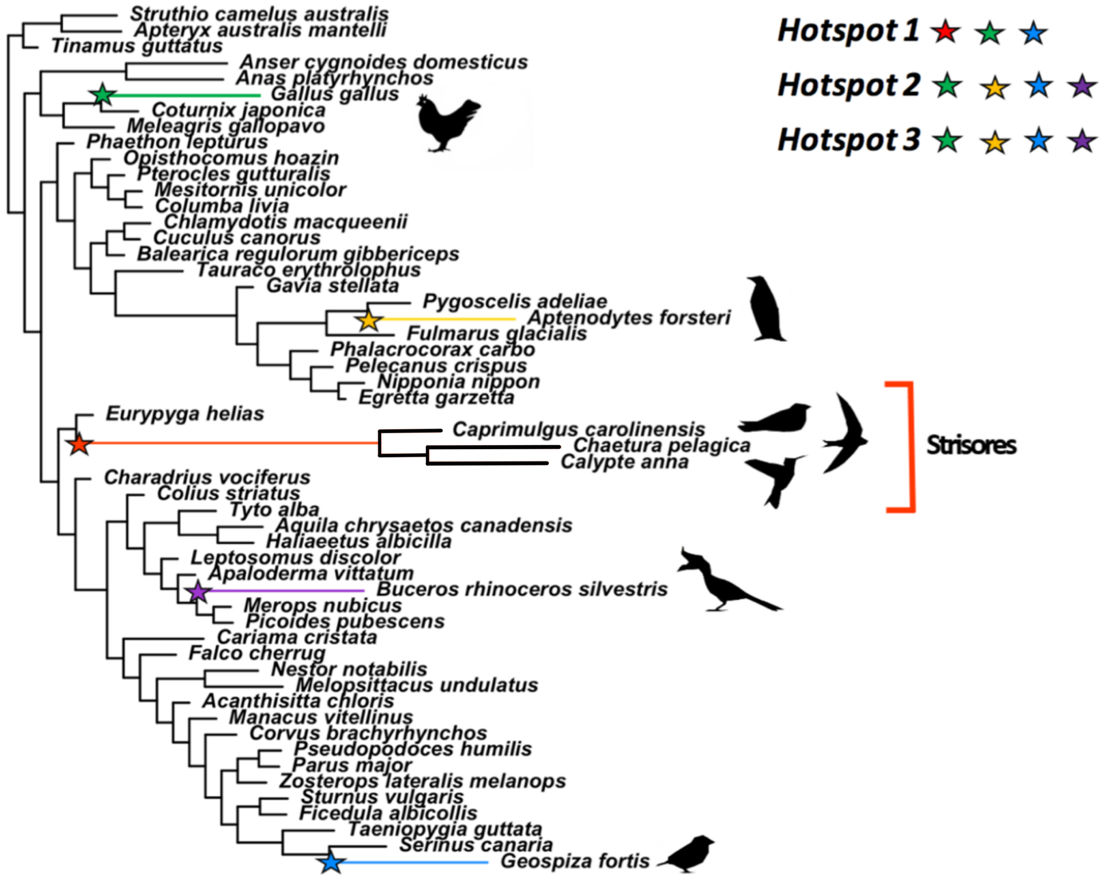

Figure S3: **An illustration of the hotspot approach containing phylogeny and the five fastest rapidly-evolving branches selected for hotspot model.** For the phylogeny, branch lengths correspond to the scaled rate of morphological beak shape evolution. Branches coloured and indicated with a star are rapidly-evolving branches that feature in the hotspot models. Because the number of available gene sequences vary per species, the fastest branches may differ for a particular gene. The key shows branches found in each hotspot model. In hotspot 1, branches found include: Strisores (consisting of nightjars and their allies, swifts and hummingbirds), Darwin's finches, and Phasianidae (represented by the red-jungle fowl). In hotspot 2, branches include: Darwin's finches, Phasianidae, Aptenodytes (represented by the emperor penguin) and Buceros (represented by the Rhinoceros hornbill). Hotspot 3 contains the same branches as hotspot 2

### Supporting Tables

#### Supplementary Table S1

Table S1: **Known candidate genes associated with beak shape morphology and size**

| *Gene symbol* | Gene name | Description |
|---|---|---|
| *ALX1* | ALX Homeobox 1 | Implicated in Lamichhaney et al (2015) as principle gene in a major locus contributing to beak shape diversity across Darwin's finches |
| *BMP2* | Bone Morphogenetic Protein 2 | Shown to correlate with beak size but not shape (Abzhanov et al, 2004). |
| *BMP4* | Bone Morphogenetic Protein 4 | Shown to correlate strongly with deep and broad beak morphology (Abzhanov et al, 2004). |
| *BMP7* | Bone Morphogenetic Protein 7 | Shown to correlate with beak size but not shape (Abzhanov et al, 2004). |
| *CALM1* | Calmodulin 1 | Shown to correlate with thin, elongated beak morphologies (Abzhanov et al, 2006). |
| *COL4A5* | Collagen Type IV Alpha 5 Chain | Shown to influence beak shape in great tits (Parus major) (Bosse et al., 2018) |
| *DKK3* | Dickkopf WNT Signaling Pathway Inhibitor 3 | Incidicated to influence different beak shapes in Darwin's finches through expression variation (Mallarino et al., 2011) |
| *DLK1* | Delta Like Non-Canonical Notch Ligand 1 | Shown in Chaves et al (2016) to correlate with beak size in Darwin's finches. |
| *FOXC1* | Forkhead Box C1 | In the largest Fst value regions between Darwin's Finches with different beak sizes (Lamichhaney et al 2015). |
| *GSC* | Goosecoid Homeobox | In the largest Fst value regions between Darwin's Finches with different beak sizes (Lamichhaney et al 2015). |
| *HMGA2* | High Mobility Group AT-Hook 2 | Implicated in Lamichhaney et al (2016) to influence beak size in species of Darwin's finches and in Chaves et al (2016). |
| *LEMD3* | LEM Domain Containing 3 | Part of a locus with significant influence on beak size In Darwin's finches (Lamichhaney et al 2016) |
| *LRRIQ1* | Leucine Rich Repeats And IQ Motif Containing 1 | Part of a locus with significant influence on beak size In Darwin's finches (Lamichhaney et al 2016) |
| *MSRB3* | Methionine Sulfoxide Reductase B3 | Part of a locus with significant influence on beak size In Darwin's finches (Lamichhaney et al 2016) |
| *RDH14* | Retinol Dehydrogenase 14 | In the largest Fst value regions between Darwin's Finches with different beak sizes (Lamichhaney et al 2015). |
| *TGFBR2* | Transforming Growth Factor Beta Receptor 2 | Incidicated to influence different beak shapes in Darwin's finches through expression variation (Mallarino et al., 2011) |
| *WIF1* | WNT Inhibitory Factor 1 | Part of a locus with significant influence on beak size In Darwin's finches (Lamichhaney et al 2016) |
| *IGF1* | Insulin-like growth factor 1 | Associated with bill size in Pyrenestes ostrinus (Vonholdt et al., 2018) |

**Supplementary Table S2**

Table S2: **Species names and file locations used for the whole genome alignment.**

| *Species name full* | Coding analysis | ASCHE analysis | Location (ftp://ftp.ncbi.nlm.nih.gov/genomes/all/GCA) and version |
|---|---|---|---|
| *Acanthisitta chloris* | x | x | /000/695/815/GCA_000695815.1_ASM69581v1/GCA_000695815.1_ASM69581v1_genomic.fna.gz |
| *Amazona aestiva* | | x | /001/420/675/GCA_001420675.1_ASM142067v1/GCA_001420675.1_ASM142067v1_genomic.fna.gz |
| *Amazona vittata* | | x | /000/332/375/GCA_000332375.1_AV1/GCA_000332375.1_AV1_genomic.fna.gz |
| *Anas platyrhynchos* | x | x | /000/355/885/GCA_000355885.1_BGI_duck_1.0/GCA_000355885.1_BGI_duck_1.0_genomic.fna.gz |
| *Anser cygnoides domesticus* | x | x | /000/971/095/GCA_000971095.1_AnsCyg_PRJNA183603_v1.0/GCA_000971095.1_AnsCyg_PRJNA183603_v1.0_genomic.fna.gz |
| *Apaloderma vittatum* | x | x | /000/703/405/GCA_000703405.1_ASM70340v1/GCA_000703405.1_ASM70340v1_genomic.fna.gz |
| *Aptenodytes forsteri* | x | x | /000/699/145/GCA_000699145.1_ASM69914v1/GCA_000699145.1_ASM69914v1_genomic.fna.gz |
| *Apteryx australis mantelli* | x | x | /001/039/765/GCA_001039765.2_AptMant0/GCA_001039765.2_AptMant0_genomic.fna.gz |
| *Aquila chrysaetos canadensis* | x | x | /000/766/835/GCA_000766835.1_Aquila_chrysaetos-1.0.2/GCA_000766835.1_Aquila_chrysaetos-1.0.2_genomic.fna.gz |
| *Ara macao* | | x | /000/400/695/GCA_000400695.1_SMACv1.1/GCA_000400695.1_SMACv1.1_genomic.fna.gz |
| *Balearica regulorum gibbericeps* | x | x | /000/709/895/GCA_000709895.1_ASM70989v1/GCA_000709895.1_ASM70989v1_genomic.fna.gz |
| *Buceros rhinoceros silvestris* | x | x | /000/710/305/GCA_000710305.1_ASM71030v1/GCA_000710305.1_ASM71030v1_genomic.fna.gz |
| *Calidris pugnax* | x | x | /001/431/845/GCA_001431845.1_ASM143184v1/GCA_001431845.1_ASM143184v1_genomic.fna.gz |
| *Calypte anna* | x | x | /000/699/085/GCA_000699085.1_ASM69908v1/GCA_000699085.1_ASM69908v1_genomic.fna.gz |
| *Caprimulgus carolinensis* | x | x | /000/700/745/GCA_000700745.1_ASM70074v1/GCA_000700745.1_ASM70074v1_genomic.fna.gz |
| *Cariama cristata* | x | x | /000/690/535/GCA_000690535.1_ASM69053v1/GCA_000690535.1_ASM69053v1_genomic.fna.gz |
| *Cathartes aura* | | x | /000/699/945/GCA_000699945.1_ASM69994v1/GCA_000699945.1_ASM69994v1_genomic.fna.gz |
| *Chaetura pelagica* | x | x | /000/747/805/GCA_000747805.1_ChaPel_1.0/GCA_000747805.1_ChaPel_1.0_genomic.fna.gz |
| *Charadrius vociferus* | x | x | /000/708/025/GCA_000708025.2_ASM70802v2/GCA_000708025.2_ASM70802v2_genomic.fna.gz |
| *Chlamydotis macqueenii* | x | x | /000/695/195/GCA_000695195.1_ASM69519v1/GCA_000695195.1_ASM69519v1_genomic.fna.gz |
| *Colinus virginianus* | | x | /000/599/465/GCA_000599465.1_NB1.1/GCA_000599465.1_NB1.1_genomic.fna.gz |
| *Colius striatus* | x | x | /000/690/715/GCA_000690715.1_ASM69071v1/GCA_000690715.1_ASM69071v1_genomic.fna.gz |
| *Columba livia* | x | x | /001/887/795/GCA_001887795.1_colLiv2/GCA_001887795.1_colLiv2_genomic.fna.gz |
| *Corvus brachyrhynchos* | x | x | /000/691/975/GCA_000691975.1_ASM69197v1/GCA_000691975.1_ASM69197v1_genomic.fna.gz |
| *Corvus cornix cornix* | x | x | /000/738/735/GCA_000738735.1_Hooded_Crow_genome/GCA_000738735.1_Hooded_Crow_genome_genomic.fna.gz |
| *Coturnix japonica* | x | x | /000/511/605/GCA_000511605.2_Coja_2.0a/GCA_000511605.2_Coja_2.0a_genomic.fna.gz |
| *Cuculus canorus* | x | x | /000/709/325/GCA_000709325.1_ASM70932v1/GCA_000709325.1_ASM70932v1_genomic.fna.gz |
| *Egretta garzetta* | x | x | /000/687/185/GCA_000687185.1_ASM68718v1/GCA_000687185.1_ASM68718v1_genomic.fna.gz |
| *Eurypyga helias* | x | x | /000/690/775/GCA_000690775.1_ASM69077v1/GCA_000690775.1_ASM69077v1_genomic.fna.gz |
| *Falco cherrug* | x | x | /000/337/975/GCA_000337975.1_F_cherrug_v1.0/GCA_000337975.1_F_cherrug_v1.0_genomic.fna.gz |
| *Falco peregrinus* | x | x | /001/887/755/GCA_001887755.1_falPer2/GCA_001887755.1_falPer2_genomic.fna.gz |
| *Ficedula albicollis* | x | x | /000/247/815/GCA_000247815.2_FicAlb1.5/GCA_000247815.2_FicAlb1.5_genomic.fna.gz |
| *Fulmarus glacialis* | x | x | /000/690/835/GCA_000690835.1_ASM69083v1/GCA_000690835.1_ASM69083v1_genomic.fna.gz |
| *Gallus gallus* | x | x | /000/002/315/GCA_000002315.3_Gallus_gallus-5.0/GCA_000002315.3_Gallus_gallus-5.0_genomic.fna.gz |
| *Gavia stellata* | x | x | /000/690/875/GCA_000690875.1_ASM69087v1/GCA_000690875.1_ASM69087v1_genomic.fna.gz |
| *Geospiza fortis* | x | x | /000/277/835/GCA_000277835.1_GeoFor_1.0/GCA_000277835.1_GeoFor_1.0_genomic.fna.gz |
| *Haliaeetus albicilla* | x | x | /000/691/405/GCA_000691405.1_ASM69140v1/GCA_000691405.1_ASM69140v1_genomic.fna.gz |
| *Haliaeetus leucocephalus* | x | x | /000/737/465/GCA_000737465.1_Haliaeetus_leucocephalus-4.0/GCA_000737465.1_Haliaeetus_leucocephalus-4.0_genomic.fna.gz |
| *Lepidothrix coronata* | | x | /001/604/755/GCA_001604755.1_Lepidothrix_coronata-1.0/GCA_001604755.1_Lepidothrix_coronata-1.0_genomic.fna.gz |
| *Leptosomus discolor* | x | x | /000/691/785/GCA_000691785.1_ASM69178v1/GCA_000691785.1_ASM69178v1_genomic.fna.gz |
| *Lyrurus tetrix tetrix* | | x | /000/586/395/GCA_000586395.1_tetTet1/GCA_000586395.1_tetTet1_genomic.fna.gz |
| *Manacus vitellinus* | x | x | /000/692/015/GCA_000692015.2_ASM69201v2/GCA_000692015.2_ASM69201v2_genomic.fna.gz |
| *Meleagris gallopavo* | x | x | /000/146/605/GCA_000146605.3_Turkey_5.0/GCA_000146605.3_Turkey_5.0_genomic.fna.gz |
| *Melopsittacus undulatus* | x | x | /000/238/935/GCA_000238935.1_Melopsittacus_undulatus_6.3/GCA_000238935.1_Melopsittacus_undulatus_6.3_genomic.fna.gz |
| *Merops nubicus* | x | x | /000/691/845/GCA_000691845.1_ASM69184v1/GCA_000691845.1_ASM69184v1_genomic.fna.gz |
| *Mesitornis unicolor* | x | x | /000/695/765/GCA_000695765.1_ASM69576v1/GCA_000695765.1_ASM69576v1_genomic.fna.gz |
| *Nestor notabilis* | x | x | /000/696/875/GCA_000696875.1_ASM69687v1/GCA_000696875.1_ASM69687v1_genomic.fna.gz |
| *Nipponia nippon* | x | x | /000/708/225/GCA_000708225.1_ASM70822v1/GCA_000708225.1_ASM70822v1_genomic.fna.gz |
| *Opisthocomus hoazin* | x | x | /000/692/075/GCA_000692075.1_ASM69207v1/GCA_000692075.1_ASM69207v1_genomic.fna.gz |
| *Parus major* | x | x | /001/522/545/GCA_001522545.2_Parus_major1.1/GCA_001522545.2_Parus_major1.1_genomic.fna.gz |
| *Passer domesticus* | | x | /001/700/915/GCA_001700915.1_Passer_domesticus-1.0/GCA_001700915.1_Passer_domesticus-1.0_genomic.fna.gz |
| *Pelecanus crispus* | x | x | /000/687/375/GCA_000687375.1_ASM68737v1/GCA_000687375.1_ASM68737v1_genomic.fna.gz |
| *Phaethon lepturus* | x | x | /000/687/285/GCA_000687285.1_ASM68728v1/GCA_000687285.1_ASM68728v1_genomic.fna.gz |

Table S2: **Species names and file locations used for the whole genome alignment.** *(continued)*

| *Species name full* | Coding analysis | ASCHE analysis | Location (ftp://ftp.ncbi.nlm.nih.gov/genomes/all/GCA) and version |
|---|---|---|---|
| *Phalacrocorax carbo* | x | x | /000/708/925/GCA_000708925.1_ASM70892v1/GCA_000708925.1_ASM70892v1_genomic.fna.gz |
| *Phoenicopterus ruber ruber* | | x | /000/687/265/GCA_000687265.1_ASM68726v1/GCA_000687265.1_ASM68726v1_genomic.fna.gz |
| *Phylloscopus plumbeitarsus* | | x | /001/655/115/GCA_001655115.1_GWplu1.0/GCA_001655115.1_GWplu1.0_genomic.fna.gz |
| *Picoides pubescens* | x | x | /000/699/005/GCA_000699005.1_ASM69900v1/GCA_000699005.1_ASM69900v1_genomic.fna.gz |
| *Podiceps cristatus* | | x | /000/699/545/GCA_000699545.1_ASM69954v1/GCA_000699545.1_ASM69954v1_genomic.fna.gz |
| *Pseudopodoces humilis* | x | x | /000/331/425/GCA_000331425.1_PseHum1.0/GCA_000331425.1_PseHum1.0_genomic.fna.gz |
| *Pterocles gutturalis* | x | x | /000/699/245/GCA_000699245.1_ASM69924v1/GCA_000699245.1_ASM69924v1_genomic.fna.gz |
| *Pygoscelis adeliae* | x | x | /000/699/105/GCA_000699105.1_ASM69910v1/GCA_000699105.1_ASM69910v1_genomic.fna.gz |
| *Serinus canaria* | x | x | /000/534/875/GCA_000534875.1_SCA1/GCA_000534875.1_SCA1_genomic.fna.gz |
| *Setophaga coronata coronata* | | x | /001/746/935/GCA_001746935.1_mywagenomev1.1/GCA_001746935.1_mywagenomev1.1_genomic.fna.gz |
| *Struthio camelus australis* | x | x | /000/698/965/GCA_000698965.1_ASM69896v1/GCA_000698965.1_ASM69896v1_genomic.fna.gz |
| *Sturnus vulgaris* | x | x | /001/447/265/GCA_001447265.1_Sturnus_vulgaris-1.0/GCA_001447265.1_Sturnus_vulgaris-1.0_genomic.fna.gz |
| *Taeniopygia guttata* | x | x | /000/151/805/GCA_000151805.2_Taeniopygia_guttata-3.2.4/GCA_000151805.2_Taeniopygia_guttata-3.2.4_genomic.fna.gz |
| *Tauraco erythrolophus* | x | x | /000/709/365/GCA_000709365.1_ASM70936v1/GCA_000709365.1_ASM70936v1_genomic.fna.gz |
| *Tinamus guttatus* | x | x | /000/705/375/GCA_000705375.2_ASM70537v2/GCA_000705375.2_ASM70537v2_genomic.fna.gz |
| *Tympanuchus cupido pinnatus* | | x | /001/870/855/GCA_001870855.1_T_cupido_pinnatus_GPC_3440_v1/GCA_001870855.1_T_cupido_pinnatus_GPC_3440_v1_genomic.fna.gz |
| *Tyto alba* | | x | /000/687/205/GCA_000687205.1_ASM68720v1/GCA_000687205.1_ASM68720v1_genomic.fna.gz |
| *Zonotrichia albicollis* | x | x | /000/385/455/GCA_000385455.1_Zonotrichia_albicollis-1.0.1/GCA_000385455.1_Zonotrichia_albicollis-1.0.1_genomic.fna.gz |
| *Zosterops lateralis melanops* | | x | /001/281/735/GCA_001281735.1_ASM128173v1/GCA_001281735.1_ASM128173v1_genomic.fna.gz |

**Supplementary Table S3**

Table S3: **Top 20 identified motifs from 39,806 genomic regions** that show significant substitution rate variation in a phylogeny-based approach were branches were binned according their beak shape morphological rate. The canonical sequences of the 20 motifs are listed along with the number of predictions from the genomic regions, the respective sequence logos and the top 5 GO predictions.

| Motif | Logo | Predictions | Top 5 specific predictions |
|---|---|---|---|
| AAAYR | 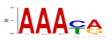 | 63 | MF olfactory receptor activity<br>BP sensory perception of smell<br>BP G-protein coupled receptor protein signaling pathway<br>BP calcium-dependent cell-cell adhesion<br>MF taste receptor activity |
| ACGT | 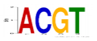 | 373 | MF RNA binding<br>BP nuclear mRNA splicing, via spliceosome<br>CC spliceosomal complex<br>BP rRNA processing<br>BP cell division |
| ACRG | 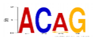 | 219 | BP G-protein coupled receptor protein signaling pathway<br>MF serine-type endopeptidase activity<br>BP defense response to bacterium<br>MF hormone activity<br>MF serine-type endopeptidase inhibitor activity |
| AWTAAW | 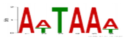 | 15 | MF olfactory receptor activity<br>BP sensory perception of smell<br>BP G-protein coupled receptor protein signaling pathway<br>BP response to stimulus<br>BP gene expression |
| AWTTAC | 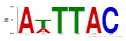 | 15 | MF olfactory receptor activity<br>BP sensory perception of smell<br>BP G-protein coupled receptor protein signaling pathway<br>BP inflammatory response<br>MF eukaryotic cell surface binding |
| BCCATTA | 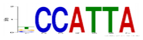 | 13 | MF olfactory receptor activity<br>BP sensory perception of smell<br>BP G-protein coupled receptor protein signaling pathway<br>BP response to stimulus<br>MF motor activity |
| CACG | 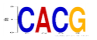 | 438 | BP rRNA processing<br>MF ATP binding<br>BP DNA repair<br>MF translation regulator activity<br>BP protein folding |
| CAG | 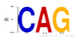 | 631 | MF calcium ion binding<br>MF serine-type endopeptidase activity<br>CC keratin filament<br>MF potassium ion binding<br>BP excretion |
| CAKCTGB | 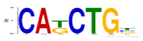 | 58 | CC extracellular space<br>BP muscle contraction<br>CC proteinaceous extracellular matrix<br>MF calcium ion binding<br>CC Z disc |
| CATAAAHC | 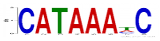 | 18 | MF olfactory receptor activity<br>BP sensory perception of smell<br>BP G-protein coupled receptor protein signaling pathway<br>BP defense response<br>BP immune response |
| CTBCC | 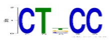 | 765 | MF potassium ion binding<br>BP potassium ion transport<br>MF protein homodimerization activity<br>MF growth factor activity<br>MF extracellular matrix structural constituent |
| CTBCWG | 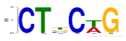 | 424 | CC extracellular space<br>CC proteinaceous extracellular matrix<br>MF calcium ion binding<br>CC keratin filament<br>MF sugar binding |
| CTCCTMC | 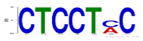 | 394 | BP transmembrane receptor protein tyrosine kinase signaling pathway<br>BP anterior/posterior pattern formation<br>BP lung development<br>BP gland development<br>MF SH3 domain binding |
| CTGKVA | 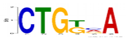 | 125 | MF serine-type endopeptidase activity<br>BP excretion<br>CC keratin filament<br>BP innate immune response<br>BP regulation of production of small RNA involved in gene silencing by RNA |
| DAAWTA | 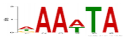 | 19 | MF olfactory receptor activity<br>BP sensory perception of smell<br>BP G-protein coupled receptor protein signaling pathway<br>BP defense response<br>CC ER to Golgi transport vesicle |
| GGGATTW | 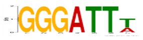 | 17 | MF olfactory receptor activity<br>BP sensory perception of smell<br>BP phototransduction<br>BP nucleobase, nucleoside, nucleotide and nucleic acid metabolic process<br>BP translation |
| GTGGGTGK | 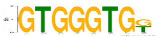 | 456 | CC integral to plasma membrane<br>BP muscle contraction<br>MF sequence-specific DNA binding<br>CC receptor complex<br>MF transcription factor activity |
| MCATATGK | 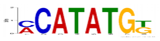 | 56 | MF olfactory receptor activity<br>BP sensory perception of smell<br>BP G-protein coupled receptor protein signaling pathway<br>BP defense response to bacterium<br>MF serine-type endopeptidase inhibitor activity |
| TTYCCW | 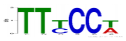 | 197 | MF olfactory receptor activity<br>BP sensory perception of smell<br>BP G-protein coupled receptor protein signaling pathway<br>CC extracellular space<br>BP signal transduction |
| WAAYGW | 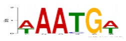 | 44 | MF olfactory receptor activity<br>BP sensory perception of smell<br>BP G-protein coupled receptor protein signaling pathway<br>MF taste receptor activity<br>BP defense response to bacterium |

#### Supplemental References

Afanasyeva A, Bockwoldt M, Cooney CR, Heiland I, Gossmann TI. 2018. Human long intrinsically disordered protein regions are frequent targets of positive selection. *Genome research* **28**: 975–982. http://www.ncbi.nlm.nih.gov/pubmed/29858274 http://www.pubmedcentral.nih.gov/articlerender.fcgi?artid=PMC6028134.

Bailey TL. 2011. DREME: motif discovery in transcription factor ChIP-seq data. *Bioinformatics* **27**: 1653–1659. https://academic.oup.com/bioinformatics/article-lookup/doi/10.1093/bioinformatics/btr261.

Benjamini Y, Hochberg Y. 1995. Controlling the false discovery rate: a practical and powerful approach to multiple testing. *Journal of the Royal Statistical Society Series B (Methodological)*.

Brunskill EW, Potter AS, Distasio A, Dexheimer P, Plassard A, Aronow BJ, Potter SS. 2014. A gene expression atlas of early craniofacial development. *Developmental biology* **391**: 133–46.

Cooney CR, Bright JA, Capp EJR, Chira AM, Hughes EC, Moody CJA, Nouri LO, Varley ZK, Thomas GH. 2017. Mega-evolutionary dynamics of the adaptive radiation of birds. *Nature* **542**: 344–347. http://www.nature.com/doifinder/10.1038/nature21074.

Fletcher W, Yang Z. 2009. INDELible: A Flexible Simulator of Biological Sequence Evolution. *Molecular Biology and Evolution* **26**: 1879–1888. https://academic.oup.com/mbe/article-lookup/doi/10.1093/molbev/msp098.

Gupta S, Stamatoyannopoulos JA, Bailey TL, Noble W. 2007. Quantifying similarity between motifs. *Genome Biology* **8**: R24. http://genomebiology.biomedcentral.com/articles/10.1186/gb-2007-8-2-r24.

Hackett SJ, Kimball RT, Reddy S, Bowie RCK, Braun EL, Braun MJ, Chojnowski JL, Cox WA, Han K-L, Harshman J, et al. 2008. A Phylogenomic Study of Birds Reveals Their Evolutionary History. *Science* **320**: 1763–1768. http://www.sciencemag.org/cgi/doi/10.1126/science.1157704.

Hillier LW, Miller W, Birney E, Warren W, Hardison RC, Ponting CP, Bork P, Burt DW, Groenen MAM, Delany ME, et al. 2004. Sequence and comparative analysis of the chicken genome provide unique perspectives on vertebrate evolution. *Nature* **432**: 695–716. http://www.nature.com/doifinder/10.1038/nature03154.

Li L, Stoeckert CJJ, Roos DS. 2003. OrthoMCL: Identification of Ortholog Groups for Eukaryotic Genomes. *Genome Research* **13**: 2178–2189. http://genome.cshlp.org/cgi/content/full/13/9/2178.

Lloyd SP. 1982. Least Squares Quantization in PCM. *IEEE Transactions on Information Theory*.

Löytynoja A, Goldman N. 2008. Phylogeny-aware gap placement prevents errors in sequence alignment and evolutionary analysis. *Science* **320**: 1632–1635.

O'Leary NA, Wright MW, Brister JR, Ciufo S, Haddad D, McVeigh R, Rajput B, Robbertse B, Smith-White B, Ako-Adjei D, et al. 2016. Reference sequence (RefSeq) database at NCBI: current status, taxonomic expansion, and functional annotation. *Nucleic Acids Research* **44**: D733–D745. https://academic.oup.com/nar/article-lookup/doi/10.1093/nar/gkv1189.

Östlund G, Schmitt T, Forslund K, Köstler T, Messina DN, Roopra S, Frings O, Sonnhammer EL. 2009. Inparanoid 7: New algorithms and tools for eukaryotic orthology analysis. *Nucleic Acids Research* **38**: 196–203.

Prum RO, Berv JS, Dornburg A, Field DJ, Townsend JP, Lemmon EM, Lemmon AR. 2015. A comprehensive phylogeny of birds (Aves) using targeted next-generation DNA sequencing. *Nature* **526**: 569–573. http://www.nature.com/doifinder/10.1038/nature15697.

R Core Team. 2018. *R: A Language and Environment for Statistical Computing*. R Foundation for Statistical Computing, Vienna, Austria https://www.r-project.org/.

Revell LJ. 2012. phytools: An R package for phylogenetic comparative biology (and other things). *Methods in Ecology and Evolution*.

Suyama M, Torrents D, Bork P. 2006. PAL2NAL: Robust conversion of protein sequence alignments into the corresponding codon alignments. *Nucleic Acids Research* **34**: 609–612.

Szklarczyk D, Franceschini A, Wyder S, Forslund K, Heller D, Huerta-Cepas J, Simonovic M, Roth A, Santos A, Tsafou KP, et al. 2015. STRING v10: Protein-protein interaction networks, integrated over the tree of life. *Nucleic Acids Research* **43**: D447–D452.

Talavera G, Castresana J. 2007. Improvement of phylogenies after removing divergent and ambiguously aligned blocks from protein sequence alignments. *Systematic Biology* **56**: 564–577.

Venditti C, Meade A, Pagel M. 2011. Multiple routes to mammalian diversity. *Nature* **479**: 393–396. http://dx.doi.org/10.1038/nature10516.

Wang J, Vasaikar S, Shi Z, Greer M, Zhang B. 2017. WebGestalt 2017: a more comprehensive, powerful, flexible and interactive gene set enrichment analysis toolkit. *Nucleic Acids Research* **45**: W130–W137. https://academic.oup.com/nar/article-lookup/doi/10.1093/nar/gkx356.

Warren WC, Hillier LW, Tomlinson C, Minx P, Kremitzki M, Graves T, Markovic C, Bouk N, Pruitt KD, Thibaud-Nissen F, et al. 2017. A New Chicken Genome Assembly Provides Insight into Avian Genome Structure. *G3 Genes|Genomes|Genetics* **7**: 109–117. http://g3journal.org/lookup/doi/10.1534/g3.116.035923.

Wilkinson L. 2011. ggplot2: Elegant Graphics for Data Analysis by WICKHAM, H. *Biometrics*.

Wu M, Chatterji S, Eisen JA. 2012. Accounting for alignment uncertainty in phylogenomics. *PLoS ONE* **7**: 1–10.

Yang Z. 1998. Likelihood ratio tests for detecting positive selection and application to primate lysozyme evolution. *Molecular Biology and Evolution* **15**: 568–573. https://academic.oup.com/mbe/article-lookup/doi/10.1093/oxfordjournals.molbev.a025957.

Yang Z. 2007. PAML 4: Phylogenetic analysis by maximum likelihood. *Molecular Biology and Evolution* **24**: 1586–1591.

Yang Z, Nielsen R, Hasegawa M. 1998. Models of Amino Acid Substitution and Applications to Mitochondrial Protein Evolution. *Mol Biol Evol* **15**: 1600–1611.

Zhang G, Li C, Li Q, Li B, Larkin DM, Lee C, Storz JF, Antunes A, Greenwold MJ, Meredith RW, et al. 2014. Comparative genomics reveals insights into avian genome evolution and adaptation. *Science* **346**: 1311–1320. http://www.sciencemag.org/cgi/doi/10.1126/science.1251385.
